# Jedi-1/MEGF12-mediated phagocytosis controls the pro-neurogenic properties of microglia in the ventricular-subventricular zone

**DOI:** 10.1101/2023.03.03.531012

**Authors:** Vivianne E. Morrison, Matthew G. Houpert, Jonathan B. Trapani, Asa A. Brockman, Philip J. Kingsley, Ketaki A. Katdare, Hillary M. Layden, Gabriela Nguena-Jones, Alexandra J. Trevisan, Kathleen A. Maguire-Zeiss, Lawrence J. Marnett, Gregory J. Bixa, Rebecca A. Ihrie, Bruce D. Carter

## Abstract

Microglia are the primary phagocytes in the central nervous system and are responsible for clearing dead cells generated during development or disease. The phagocytic process shapes the phenotype of the microglia, which affects the local environment. A unique population of microglia reside in the ventricular-subventricular zone (V-SVZ) of neonatal mice, but how they influence this neurogenic niche is not well-understood. Here, we demonstrate that phagocytosis creates a pro-neurogenic microglial phenotype in the V-SVZ and that these microglia phagocytose apoptotic cells via the engulfment receptor Jedi-1. Deletion of Jedi-1 decreases apoptotic cell clearance, triggering the development of a neuroinflammatory phenotype, reminiscent of neurodegenerative and-age-associated microglia, that reduces neural precursor proliferation via elevated interleukin (IL)-1β signaling; inhibition of IL-1 receptor rescues precursor proliferation in vivo. Together, these results reveal a critical role for Jedi-1 in connecting microglial phagocytic activity to a phenotype that promotes neurogenesis in the developing V-SVZ.

**Graphical Abstract. Jedi-1-dependent phagocytosis supports neurogenesis via suppression of microglial inflammatory pathway activation:** Top: Wild-type Proliferative-zone-Associated Microglia (PAMs) (cyan) use the engulfment receptor Jedi-1 (‘Jedi’) to engulf apoptotic cells (yellow) in the neurogenic ventricular-subventricular zone (V-SVZ) of the early postnatal brain. Jedi activation supports neural precursor cell (NPC) proliferation and the generation of new neurons.

Bottom: Deletion of Jedi reduces microglial phagocytosis and transforms PAMs into Disease-associated Inflammatory Microglia (DIMs) characterized by the upregulation of canonical inflammatory genes and core DIM markers iden ified in the aging and neurodegenerative brain (Nlrp3, NLR family pyrin domain-containing 3; Tnf, tumor necrosis factor; Ccl4, C-C chemokine ligand 4 (also called macrophage inflammatory protein 1β); Ccr5, C-C chemokine receptor type 5). Increased interleukin-1β (IL-1β) synthesis, release, and signaling in the Jedi-null V-SVZ reduces NPC proliferation and newborn neuron number.

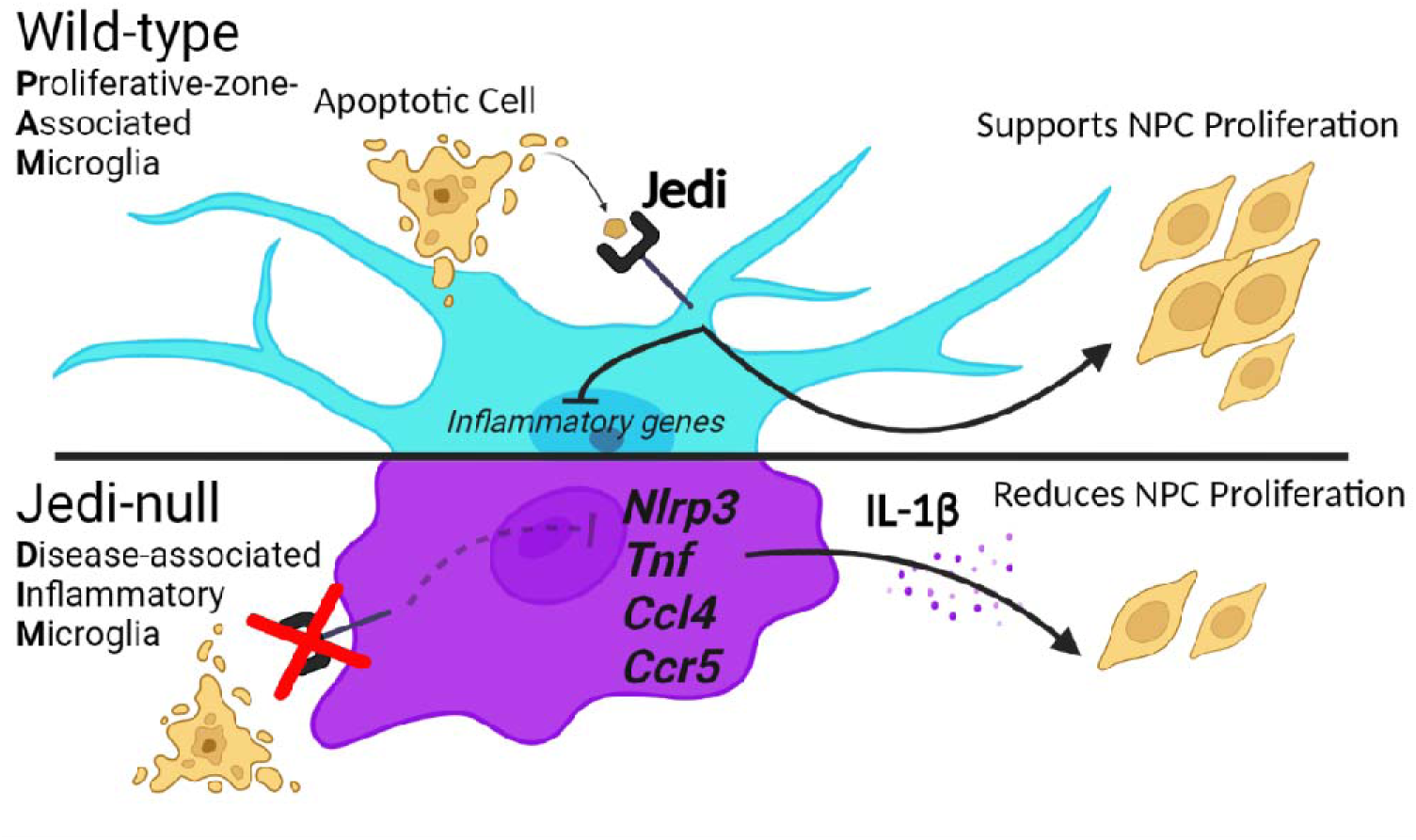

**Highlights:**

- The engulfment receptor Jedi-1 is expressed by microglia in the neonatal ventricular-subventricular zone (V-SVZ) neurogenic niche.
- *Jedi-1* knockout microglia have decreased engulfment ability, resulting in accumulation of dead cells in the V-SVZ.
- Loss of *Jedi-1* leads to a neuroinflammatory phenotype in microglia that is characteristic of neurodegenerative and age-associated microglia.
- Microglial-specific loss of *Jedi*-1 reduces neurogenesis, which is rescued *in vivo* by inhibition of interleukin-1β signaling.

## Introduction

Microglial phagocytosis is critical for sculpting the developing central nervous system (CNS) and eliminating inflammatory debris such as dead cells, myelin debris, and protein aggregates in homeostatic and disease states^1^. Phagocytosis also profoundly affects the formation of context-specific microglial identities, reflected in unique transcriptomic, morphological, and functional features. For example, after demyelinating injury and in Alzheimer’s Disease mouse models, cellular debris activates the engulfment receptor TREM2 (triggering receptor expressed on myeloid cells 2), which leads to a microglial phenotype characterized by ameboid morphology, increased proliferation and chemotaxis, and upregulated lipid metabolism, phagocytic, and lysosomal pathways^2,3^. The genetic signature of TREM2-dependent microglia is distinct from that of microglia under homeostatic conditions and is found to lesser or greater extents in varying disease contexts^3,4^. If the TREM2-dependent phenotype fails to form, then neurodegeneration progresses, and this suggests that TREM2 activity and the microglial phenotype it engenders are neuroprotective^3^. Interestingly, a transcriptomically similar genetic signature is also present in a microglial population found specifically in the ventricular-subventricular zone (V-SVZ) neurogenic niche at 1-2 weeks after birth in several animal models, including mice^5–7^. To what extent and by what mechanism these microglia perform phagocytosis, whether it plays a role in their distinctive phenotype, and whether it regulates neurogenesis are questions that have yet to be addressed.

Neurogenesis, that is, the production of neural precursor cells (NPCs) and immature neurons (neuroblasts) in the V-SVZ is robust, but many of the newborn cells undergo apoptosis as part of a normal culling process necessary for the maintenance of tissue organization and to prevent excessive cell numbers^8,9^. It has been calculated that about 30,000 NPCs are produced every day in the adult mouse V-SVZ^10^, and it is estimated that as many as 60% of those cells undergo apoptosis^11–14^. Given that the rate of neurogenesis is higher in younger animals, this results in a substantial amount of cellular debris to be cleared from the neonatal V-SVZ, and evidence suggests that V-SVZ-resident microglia engulf these apoptotic NPCs^15^. Disruption of microglial function in vivo, though not specifically of phagocytosis, negatively impacts NPC proliferation, neuronal differentiation, and survival, suggesting that neonatal V-SVZ-resident microglia are pro-neurogenic and support the neurogenic program^15–17^. Whether their pro-neurogenic functions are related to their phagocytic functions remains to be determined.

Although the neonatal V-SVZ is a unique niche, containing specialized microglia, studies in the adult neurogenic regions, including the hippocampal subgranular zone (SGZ), suggest that microglial phagocytosis can influence neurogenesis. For example, in the adult mouse, deletion of Axl and MerTK, members of the TAM (Tyro3/Axl/MerTK) family of engulfment receptors, led to (1) the accumulation of dead cells in the V-SVZ and along the path to the olfactory bulb (the principal destination of neuroblasts born in the V-SVZ), (2) the development of ameboid microglial morphology, and surprisingly, (3) an increased number of newborn neurons in the olfactory bulb^18^. In contrast, in the adult SGZ, the deletion of the engulfment receptors P2Y12 or Axl and MerTK reduced NPC proliferation and neuronal differentiation^19^. Transcriptomic profiling of phagocytic microglia in vitro suggested that widespread changes in the secretome of these cells accounted for the pro-neurogenic influence of phagocytic microglia in the adult SGZ^19^. Taken together, these results highlight an intimate relationship between microglial phagocytic ability and adult neurogenesis, though the exact nature of the relationship and the underlying mechanisms remain to be determined in both the adult and neonatal brain. Furthermore, whether the phenotype of the V-SVZ-resident microglia population is influenced directly by the phagocytic process has not been explored.

Importantly, microglia phenotypes exist in many different states across space, time, and condition, making it difficult to concisely capture the diversity and specificity of microglial states^20^. For the sake of brevity, we use several common acronyms to describe groups of cells with features that are distinct from those of microglia found in homeostatic conditions (i.e., PAMs, DAMs, and DIMs, defined below). However, it is imperative to recognize that these abbreviations are used solely to reduce the linguistic complexity that would otherwise be required to accurately delineate between microglial phenotypes and thus, do not reflect the true nature of microglial states in the brain.

Microglia found in the neonatal V-SVZ (also called ‘proliferative region-associated microglia’ (PAMs)^7^) and those that appear in response to disease (referred to as ‘disease-associated microglia’ (DAMs)^3,4^) appear to be pro-neurogenic and neuroprotective, respectively. On the other hand, another recently described population of ‘disease inflammatory microglia’ (DIMs) was found in neurodegenerative conditions and in the aged brain. Due to their expression of pro-inflammatory genes, DIMs likely contribute significantly to neurodegeneration and functional decline^5,6^. Previous literature has demonstrated a connection between neurodegeneration and dysfunctional engulfment or impaired degradation of engulfed material, suggesting that phagocytosis is a protective measure that restricts inflammatory activation of microglia^21–23^.

Overall, these findings emphasize the idea that phagocytosis is the lens through which microglia see their environment and helps to transform microglial identity to match specific contexts. To explore the relationship between phagocytosis, microglial phenotype, and neurogenesis, we investigated the presence and function of the engulfment receptor Jedi-1 in the V-SVZ. Jedi-1 (hereafter referred to as ‘Jedi’), also called Multiple EGF-like-domains 12 (MEGF12) and Platelet-Endothelial Aggregation Receptor 1 (PEAR1), is a mammalian homolog of the engulfment receptors Draper and CED-1 in flies and nematodes, respectively^24^.Characterization of Jedi in the developing peripheral nervous system (PNS) revealed its role in promoting the clearance of apoptotic sensory neurons by adjacent satellite glial precursors and its usage of the same intracellular signaling motifs employed by TREM2-DAP12 signaling^24–26^., however, the role of Jedi in the CNS is unknown.

We show that Jedi is expressed by microglia in the V-SVZ of neonatal mice. While the neonatal period extends from postnatal day 0 (P0) to P28^27^, we explicitly examined mice at P7, when postnatal NPC turnover and PAM density reach their apices^7,15,28^. Microglia in Jedi-null animals have reduced phagocytic ability, resulting in the accumulation of apoptotic cells in the V-SVZ. Moreover, Jedi-deficient microglia dramatically change their morphology, proliferative capacity, and transcriptomic profile, adopting a DIM-like phenotype. These DIM-like neonatal microglia suppress V-SVZ neurogenesis, and this reduction is rescued by blocking signaling through interleukin-1 (IL-1), a canonical inflammatory pathway. These findings establish, for the first time, a mechanistic connection between microglial phagocytic activity and the unique microglial phenotype that ensures neurogenesis in the neonatal V-SVZ. Critically, we demonstrate that Jedi is a fulcrum that controls the balance of inflammatory and anti-inflammatory pathways and defines the pro-neurogenic identity of microglia.

## Results

### Jedi is expressed in microglia in the neonatal V-SVZ

To determine whether Jedi plays a role in apoptotic cell clearance in the CNS, we assessed its protein expression in microglia in the wild-type (WT) neonatal mouse brain. Several RNA sequencing studies have shown that Jedi transcripts are present in neonatal microglia^5,7,29^. We hypothesized that Jedi would be expressed in areas with ongoing cell death, substantial debris to be cleared, and a high density of microglia that are likely phagocytic (i.e., the V-SVZ at postnatal day 7 (P7)). Using immunofluorescent labeling of IBA1 to identify microglia, we found that Jedi was expressed by a subset of the IBA1^+^ population in the V-SVZ (Figure 1A, arrows indicate Jedi^+^ (yellow) and Jedi^-^ (magenta) microglia). We also found Jedi expressed in microglia in the corpus callosum (CC) dorsal to the V-SVZ (Figure 1A, CC is visible in the upper right corner of the image, above the V-SVZ, which is identified by its high DAPI^+^ cell density). We did not detect Jedi protein in cortical or striatal microglia (Figure 1B and 1C, respectively), suggesting that Jedi expression is restricted to V-SVZ-resident microglia. This restricted expression profile may explain why the available RNA sequencing datasets, which use microglia isolated from the whole brain and not the micro-dissected V-SVZ, show only low levels of Jedi expression. Cortical and striatal microglia had appreciably less intense IBA1 staining than that seen in V-SVZ-and CC-resident microglia. Moreover, cortical and striatal microglia had more branches and finer processes than V-SVZ-and CC-resident microglia, the latter being amorphous and less morphologically complex. Increased IBA1 expression and an ameboid morphology are typically associated with a departure from homeostatic microglial states^30^, and our observations align with those of others who have noted morphological differences and varying degrees of surface-antigen staining intensity between microglia in the V-SVZ or the developing white matter and those that lie outside these zones^7,15^. Jedi expression was also detected on blood vessels (Figures 1A–1C), as has been reported in peripheral tissues^31^.

**Figure 1.**
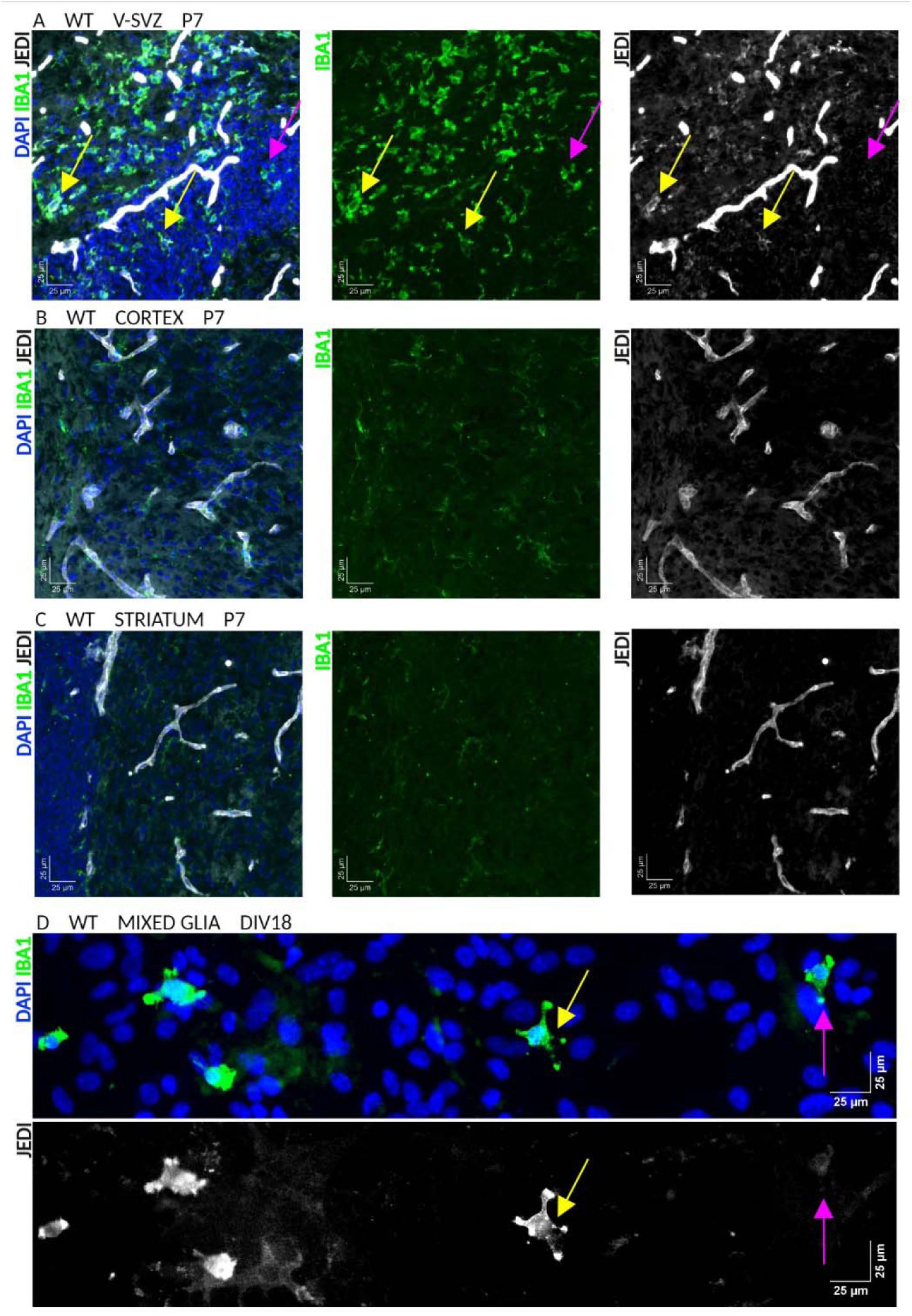
Jedi expression is present in microglia in the early postnatal V-SVZ and in vitro. 1A. Wild-type (WT) ventricular-subventricular zone (V-SVZ) at postnatal day 7 (P7). Left: Merge of DAPI (blue), IBA1 (green), and Jedi (white). Middle: IBA1 alone. Right: Jedi alone. Yellow arrows indicate IBA1+Jedi+ microglia. Magenta arrow indicates IBA1+Jedi-microglia. Long white structures are blood vessels. Scale bar: 25 microns (um). 1B. WT cortex at P7. Left: Merge of DAPI (blue), IBA1 (green), and Jedi (white). Middle: IBA1 alone. No IBA1+Jedi+ microglia were observed. Long white structures are blood vessels. Scale bar: 25um. 1C. WT striatum at P7. Left: Merge of DAPI (blue), IBA1 (green), and Jedi (white). Middle: IBA1 alone. No IBA1+Jedi+ microglia were observed. Long white structures are blood vessels. Scale bar: 25um. 1D. WT mixed glia cultures (days in vitro 18, DIV18) isolated from P7 mice. Top: Merge of DAPI (blue) and IBA1 (green). Bottom: Jedi (white) alone. Yellow arrow indicates one of several IBA1+Jedi+ microglia. Magenta arrow indicates IBA1+Jedi-microglia. Scale bar: 25um

### Cultured microglia express Jedi and Jedi is necessary for microglial phagocytosis in vitro

Immunofluorescent labeling of Jedi and IBA1 in mixed glia cultures isolated from the cerebrum showed that Jedi expression persists in culture, and like in vivo, some IBA1^+^ microglia were Jedi^+^ while others were not (Figure 1D, yellow and magenta arrowheads, respectively). A recent study that used RNA sequencing of cultured microglia showed that microglia derived from P5 wildtype mice express Jedi^32^. To focus on microglia, we isolated microglia via the shake-off method and found purity of the cultures >95%, based on immunolabeling for IBA1 and GFAP, a marker of astrocytes (Supplementary Figure 1).

To determine whether Jedi is needed for microglial phagocytosis, we used an in vitro engulfment assay using microglia cultures isolated from P7 WT and global germline Jedi knockout (JKO) mice. The assay uses carboxylated latex beads that are highly negatively charged, similar to phosphatidylserine, an established “eat-me” signal^33^. The beads are two microns in diameter, and thus, are too large to be efficiently engulfed via non-specific, non-receptor-mediated endocytic pathways (i.e., fluid-phase endocytosis, also called pinocytosis)^34^. Microglia were exposed to the beads for up to 4 hours, then the cells were very gently washed with PBS only once to remove the media without disturbing beads that were attached to the cells but not engulfed; this resulted in the retention of many unengulfed beads on the slide, as well (Figure 2A and 2B). The cells were stained with phalloidin to label filamentous actin and allow visualization of the cell soma and processes, and imaged using confocal microscopy. In the XY plane, we saw beads associated with both WT and JKO microglia (Figure 2A and 2B), and the number of beads that were associated with cells was visibly reduced in Jedi-deficient cultures. Using orthogonal views of z-stacks to differentiate between engulfed, attached, and free beads (Figure 2C; green crosshairs, yellow asterisk, and magenta arrowhead, respectively), we found a significant decrease in the proportion of microglia that were performing bead engulfment in the JKO condition relative to WT at both 2 and 4 hours after addition of the beads (Figure 2D). The phagocytic index (PI), which is calculated using the number of engulfed beads, the number of cells engulfing beads, and the total number of cells in a field of view, showed a time-dependent increase in bead engulfment in both genotypes, pointing to the likely presence of Jedi-independent engulfment pathways, but the PI of JKO microglia was significantly reduced relative to WT cells at both 2 and 4 hours (Figure 2E). A modified PI equation was used to quantify the number of beads that were attached to the cell surface but not engulfed (“Bead Binding Index”), and we found that it too was decreased in JKO microglia at both time points (Supplementary Figure 2), suggesting a defect in bead binding and/or bead retention in the absence of Jedi.

**Figure 2.**
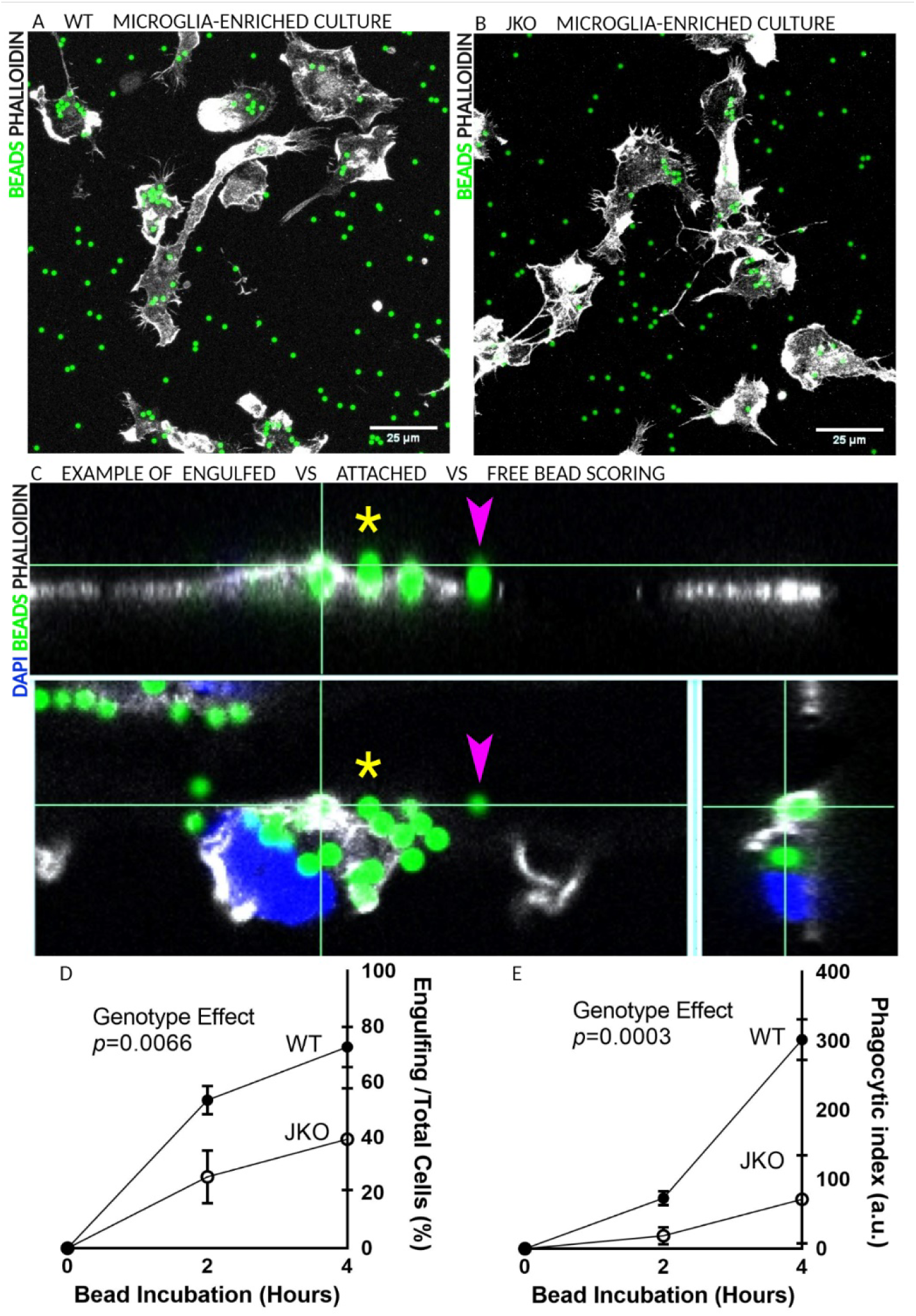
Jedi is necessary for microglial phagocytosis in vitro. 2A. Wild-type (WT) microglia stained with phalloidin (white) engulfing latex beads (green). Scale bar: 25 microns (um). 2B. Jedi knockout (JKO) microglia stained with phalloidin (white) engulfing latex beads (green). Scale bar: 25um. 2C. Example of the use of orthogonal views of a microglia (phalloidin, in white) incubated with beads (green). Microglia nucleus is stained by DAPI (blue). Top: y,z view of an engulfing cell. Green crosshairs indicate an engulfed bead. Yellow asterisk indicates an attached, but not engulfed bead. Magenta arrowhead indicates a free bead. Bottom left: x, y view of the engulfing cell. Bottom right: x, z view of the engulfing cell; only the engulfed bead is visible in this plane at this position. 2D. Quantification of the percentage of microglial cells that are engulfing beads in JKO (open circle) and WT (closed circle). Ordinary Two-way ANOVA: Column Factor (Genotype), p=0.0066; Row Factor (Time point), p=<0.0001); Interaction, p=0.0961. Each data point is the average of 3 independent microglial cultures and engulfment assays. 2E. Quantification of the phagocytic index (defined as (the number of engulfed beads/the total number of cells) x (the number of engulfing/the total number of cells) x 100) in JKO (open circle) and WT (closed circle). Ordinary Two-way ANOVA: Column Factor (Genotype), p=0.0003; Row Factor (Time point), p=<0.0001); Interaction, p=0.001. Each data point is the average of 3 independent microglial cultures and engulfment assays. Arbitrary units, a.u.

### Jedi deficiency leads to the accumulation of dead cells in the V-SVZ

We reasoned that reduced engulfment by Jedi-deficient microglia would cause dead cells to accumulate in the V-SVZ (Figure 3A, the V-SVZ is outlined). To test the hypothesis that the number of dead cells present in the V-SVZ would be higher in JKO than in WT, we used the TUNEL assay to mark cleaved DNA, the latter being an obligate step in the apoptotic process. We found that numerous TUNEL^+^ cells were present in the WT V-SVZ, consistent with the increased cell turnover that occurs in this region at P7, but the JKO V-SVZ contained significantly more TUNEL^+^ cells than the WT V-SVZ (Figure 3B–D). We found little to no cell death in the cortex of either JKO or WT mice (Figure 3E and 3G (left-most box-and-whisker plot)). In the striatum immediately adjacent to the V-SVZ, the average number of TUNEL^+^ cells per mm2 in the WT and JKO were 464.2 and 670.2, respectively (p=0.05). However, the effect size was small (206.0 ± 110.5) and the 95% confidence interval (CI) included zero (−48.71 to 460.7) (Figure 3F and 3G (right-most box-and-whisker plot)), weakening the notion that there are reproducible, physiologically meaningful changes in cell death in the striatum of JKO mice, and supporting the idea of a V-SVZ-specific effect of Jedi deficiency.

**Figure 3.**
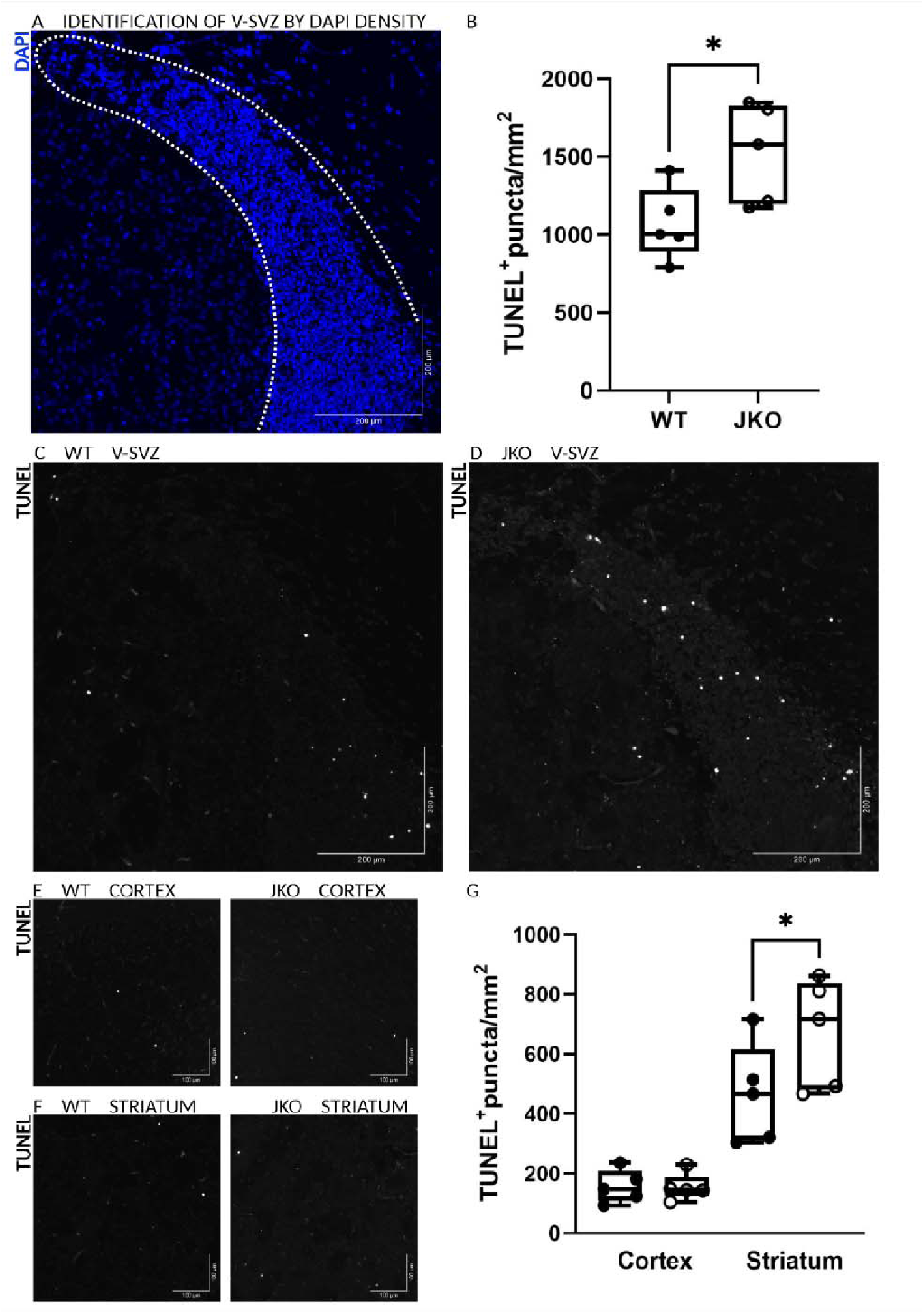
Dead cells accumulate in the V-SVZ of Jedi-deficient mice. 3A. The ventricular-subventricular zone (V-SVZ) has high DAPI signal (blue) due to high cell density. The V-SVZ is outlined by the white-dotted line. Image right side: medial. Left side: lateral. Top edge: dorsal. Bottom edge: ventral. Scale bar: 200 microns (um). 3B. Quantification of dead cells labeled by the TUNEL assay (Terminal deoxynucleotidyl transferase dUTP nick end labeling) per unit area in the V-SVZ of Jedi knockout (JKO) and wild-type (WT) mice. Unpaired one-tailed t test, * p=0.0163. 3C. WT V-SVZ has a few TUNEL+ dead cells (white). Scale bar: 200um. 3D. JKO V-SVZ has many TUNEL+ dead cells (white). Scale bar: 200um. 3E. Dead cells in the cortex. Left: WT. Right: JKO. 3F. Dead cells in the striatum. Left: WT. Right: JKO. 3G. Quantification of dead cells labeled by the TUNEL assay in the cortex and striatum of WT and JKO mice. Unpaired two-tailed t test, * p=0.0496; 95% confidence interval, −48.71 to 460.7.

### Jedi deficiency reduces phagocytosis of dead cells and increases the number of microglia and IBA1-stained area in the V-SVZ

To test whether dead cell engulfment was reduced in the JKO V-SVZ in vivo, we generated 3D reconstructions of V-SVZ-resident microglia in sections labeled with IBA1 and TUNEL. All TUNEL^+^ cells were scored as ‘engulfed’ (i.e., inside a microglial cell body or phagocytic cup; yellow asterisk in Figure 4C) or ‘non-engulfed’ (i.e., not associated with a microglial cell; magenta arrowhead in Figure 4C). Quantification revealed that a smaller proportion of the apoptotic cells in the JKO V-SVZ were engulfed relative to in WT V-SVZ (Figure 4A,4B, and 4D). While this approach cannot address whether there is increased cell death in the JKO V-SVZ, the number of microglia (Figure 4E–4G) and total IBA1^+^ area in the V-SVZ (Figure 4H–4J) were higher in JKO than in WT, and the increased number of non-engulfed apoptotic cells, even in the presence of more microglia, points to defective phagocytosis as a major contributor to the accumulation of dead cells. Our in vitro and in vivo findings support the idea that Jedi expression by microglia is necessary for the clearance of apoptotic cells in the neonatal V-SVZ.

**Figure 4.**
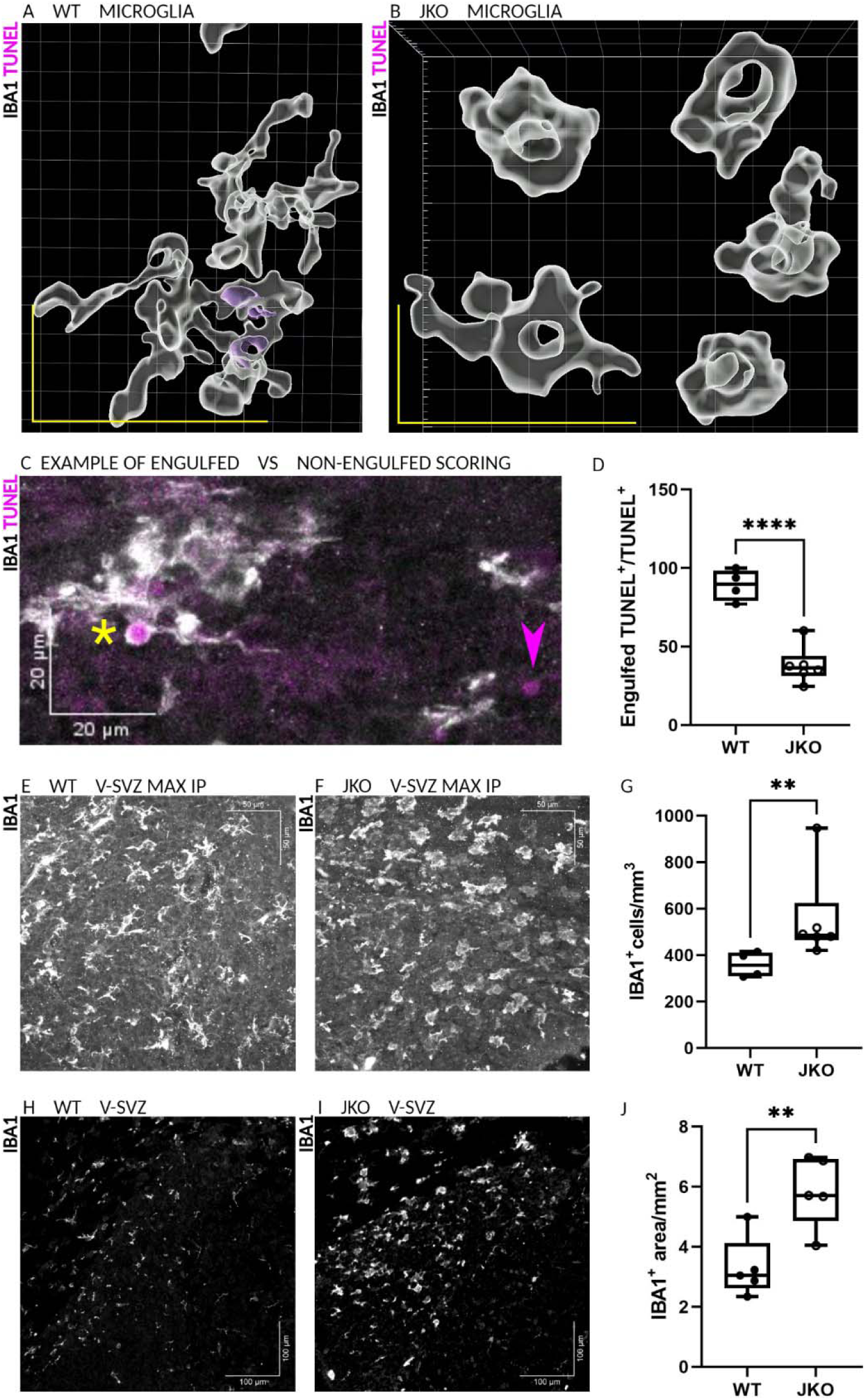
Jedi deficiency reduces phagocytosis of dead cells and increases the number of microglia and IBA1-stained area in the V-SVZ. 4A. 3D rendering of wild-type (WT) microglia (IBA1+, white) engulfing dead cells (TUNEL+, magenta) in the postnatal day 7 (P7) ventricular-subventricular zone (V-SVZ). 4B. 3D rendering of Jedi knockout (JKO) microglia (IBA1+, white) in the postnatal day 7 (P7) ventricular-subventricular zone (V-SVZ). No engulfed cells are visible in this image. 4C. Maximum intensity projection (MAX IP) showing an example of in vivo engulfment scoring. Microglia (IBA1+, white). TUNEL+ dead cells (magenta). Yellow asterisk indicates an engulfed cell (immediately adjacent on the right side). Magenta arrowhead indicates a non-engulfed dead cell. Scale bar: 20 microns (um). 4D. Quantification of the percentage of TUNEL+ cells that were contained within a microglial cell in the JKO and WT V-SVZ. Unpaired one-tailed t test, *** p<0.0001. 4E. MAX IP of IBA1 staining (white) in the P7 WT V-SVZ. Scale bar: 50um. 4F. MAX IP of IBA1 staining (white) in the P7 JKO V-SVZ. Scale bar: 50um. 4G. Quantification of the number of microglia per unit area in the V-SVZ of WT and JKO mice. Normality tests: JKO did not pass (Shapiro-Wilk test, W=0.6500, p=0.0018; Kolmogorov-Smirnov test, KS distance=0.4122, p=0.0019). Mann-Whitney test, p=0.0095. 4H. IBA1 staining (white) in WT V-SVZ at P7. Scale bar: 100um. 4I. IBA1 staining (white) in JKO V-SVZ P7. Scale bar: 100um. 4J. Quantification of IBA1+ area per unit area in the V-SVZ. Unpaired two-tailed t test, ** p=0.0062.

### Jedi-deficient microglia adopt an activated phenotype characterized by increased proliferation and ameboid morphology

While investigating TUNEL^+^ and IBA1^+^ cell numbers, we noted remarkable differences in proliferation and morphology between JKO and WT microglia, which could reflect a shift toward an inflammatory state. In non-homeostatic conditions, microglia are highly proliferative, and proliferative microglia, which were identified by incorporation of the synthetic nucleotide EdU into newly synthesized DNA, were more abundant in the V-SVZ of JKO mice relative to WT mice, which could explain the increased size of the microglial population in the Jedi-null V-SVZ (Figure 5A–5C, yellow arrowheads indicate EdU^+^IBA1^+^ cells).

**Figure 5.**
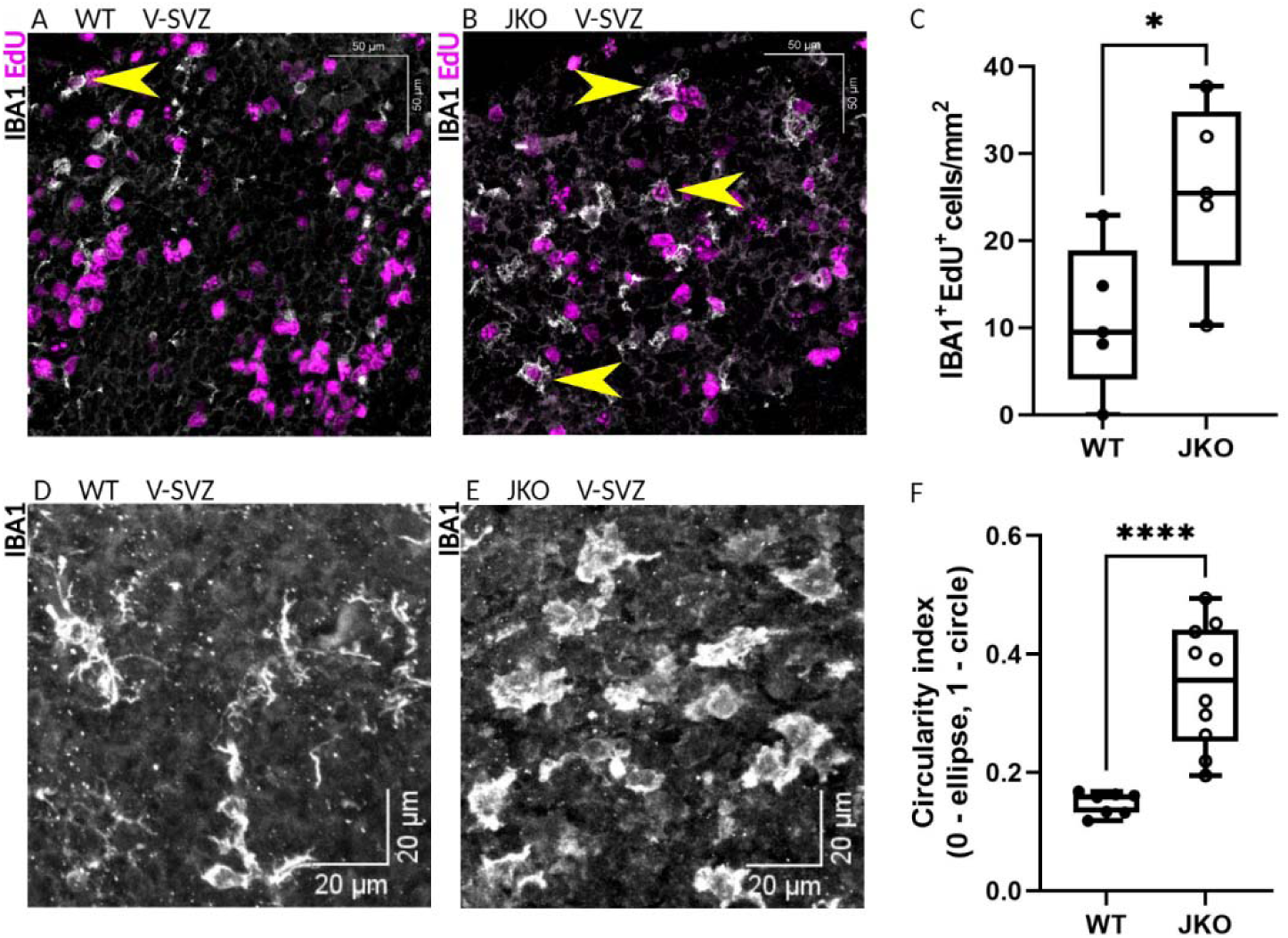
Jedi-deficient microglia adopt an activated phenotype characterized by increased proliferation and ameboid morphology. 5A. Microglia (IBA1, white) and newly divided cells (EdU (5-ethynyl-2’-deoxyuridine), magenta) in the wild-type (WT) ventricular-subventricular zone (V-SVZ) at postnatal day 7 (P7). Yellow arrowhead indicates an IBA1+EdU+ cell. Scale bar: 50 microns (um). 5B. Microglia (IBA1, white) and newly divided cells (EdU, magenta) in the Jedi knockout (JKO) V-SVZ at P7. Yellow arrowheads indicate IBA1+EdU+ cells. Scale bar: 50um. 5C. Quantification of IBA1+EdU+ cells per V-SVZ area. Unpaired one-tailed t test, * p=0.0190. 5D. Microglia (IBA1, white) in the WT V-SVZ at P7. Scale bar: 20um. 5E. Microglia (IBA1, white) in the JKO V-SVZ at P7. Scale bar: 20um. 5F. Quantification of the circularity index (4π[area]/[perimeter]2). Unpaired one-tailed t test, **** p<0.0001.

Microglial morphology generally correlates with state changes, and non-homeostatic microglia are rounder (‘ameboid’) and have fewer processes than homeostatic microglia^35^. WT microglia were elongated and had many fine branches, but JKO microglia were ameboid and had few branches (Figure 5D and 5E). We quantitatively assessed these differences using the circularity index, where 1.0 indicates a perfect circle^36^. JKO microglia had higher index values than WT microglia, reflecting their ameboid shape (Figure 5F). The increased proliferation and ameboid morphology of JKO microglia suggest that Jedi regulates pathways typically activated upon loss of brain homeostasis. This supports the idea that Jedi-dependent microglial phagocytosis plays a role in establishing and maintaining the unique phenotype of microglia in the neonatal V-SVZ.

### RNA sequencing of WT telencephalic microglia replicates the transcriptomic differences previously identified between V-SVZ and cortical microglia

To assess changes in the genetic signature of Jedi-deficient microglia, we performed bulk RNA sequencing on acutely isolated microglia from the V-SVZ and, separately, the cortex of WT and JKO animals at P7. Microglia were collected using immunomagnetic sorting based on CD11b expression (Figure 6A). Immunolabeling of IBA1 in the selected cells revealed the population to be more than 99% microglia (Supplementary Figure 3). Within the WT group, cortical and V-SVZ microglia had very different transcriptomic signatures (Figure 6B). We made two additional observations that are consistent with previous RNA sequencing data on neonatal microglia: first, we found no evidence of differential expression at P7 of so-called ‘adult homeostatic genes’ (i.e., Tmem119, P2ry12, Siglech, Selplg, Cx3cr1, Txnip, Lgmn, or Serinc3, etc.). This is consistent with the idea that neonatal expression of these genes is low, until their expression increases in juvenile microglia, and after P30, their levels are maintained in the adult and aging animal^2,4,5,7^. Second, we observed higher levels of PAM/DAM genes (i.e., Itgax, Igf1, Spp1, Gpnmb, Mamdc2, Apoc1, Cd9, Ctsb, Apoe, and Lpl) in microglia from the V-SVZ when compared to those of the cortex (Figure 6C, rose text and data points). Together, these findings confirm the presence of and our ability to detect the V-SVZ microglial population as it has been described in previous datasets.

**Figure 6.**
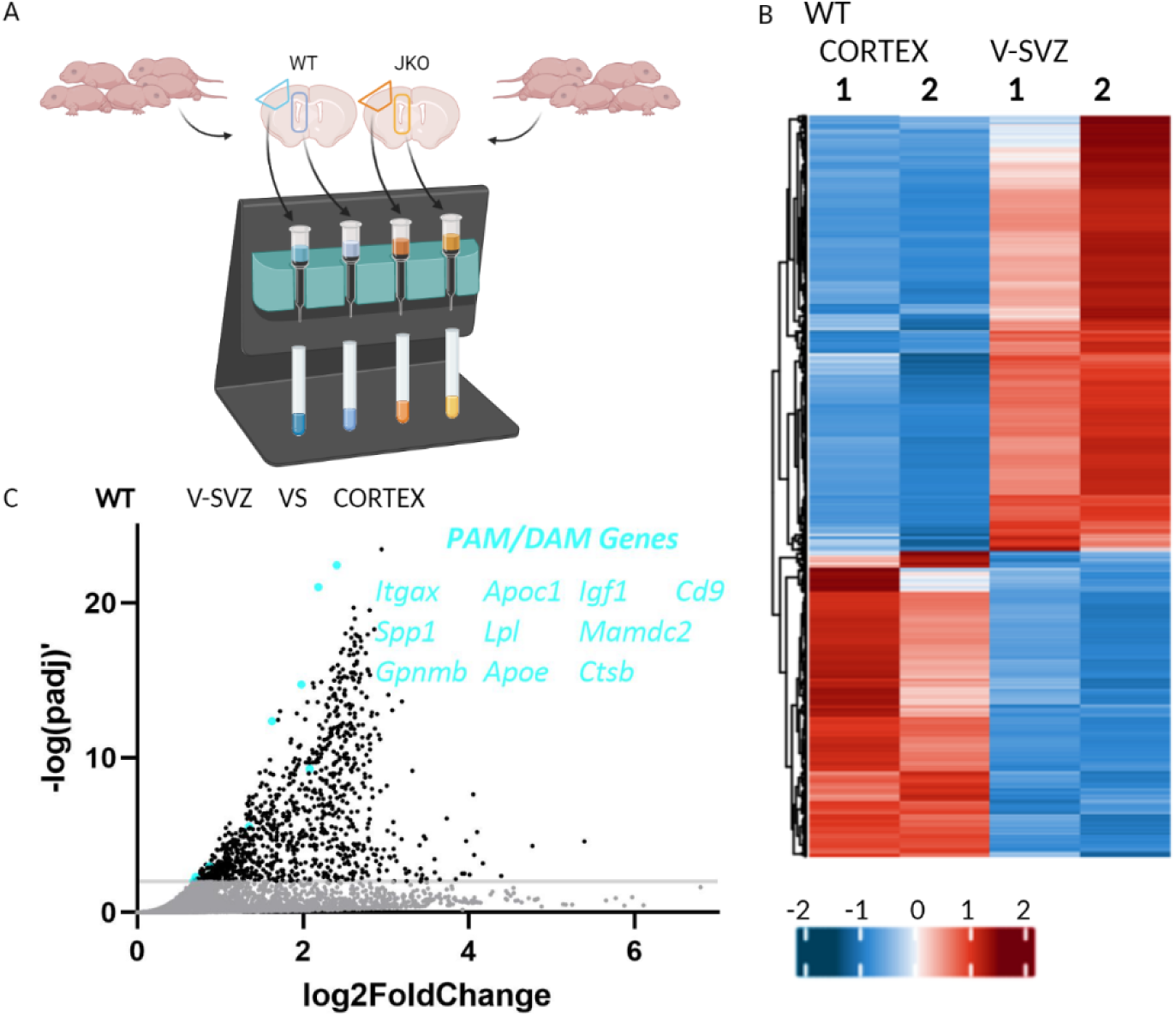
RNA sequencing confirms previously described transcriptomic differences between PAMs and cortical microglia. 6A. Workflow schematic of cortex and ventricular-subventricular zone (V-SVZ) isolation from pooled wild-type (WT) and pooled Jedi knockout (JKO) pups for magnetic cell sorting of CD11b+ microglia and bulk RNA sequencing of the collected microglia. 6B. Heatmap demonstrating clear differences in the genetic signature of cortical and V-SVZ microglia. Colors represent direction of change (blue: relative downregulation, red: relative upregulation) and degree of change (z-scores). 6C. Volcano plot illustrating the upregulation of ten core markers (rose) of proliferative-zone-associated microglia (PAM) and disease-associated microglia (DAM).

### RNA sequencing confirms defective phagocytosis of neurogenic cells by Jedi-deficient microglia

Next, we focused on differences between WT and JKO, specifically in microglia derived from the V-SVZ. Jedi knockout resulted in the upregulation of 238 genes and the downregulation of 842 genes in V-SVZ microglia relative to those in WT samples (threshold of significance set at padj < 0.01) (Figure 7A and 7B). As a result of its germline global deletion, Jedi expression was significantly reduced in JKO microglia (Figure 7B, red text and data point). Between JKO and WT cells, we found that the most differentially represented genes were associated with NPCs, neuroblasts, and oligodendrocyte lineage cells (OLCs) (i.e., Egfr, Pax6, Meis2, Insm1, Dlx1/2, Dcx, Olig2, Cspg4, Pdgfra, Sox10, Myt1, etc.), and all were reduced in JKO microglia (Figure 7C; NPC genes, blue text and data points; neuroblast genes, yellow text and data points; OLC genes, green text and data points; astrocyte genes, magenta text and data points). This is consistent with previous findings showing that microglia become enriched for genes associated with neurogenesis following phagocytosis of apoptotic neurons^19^ and that PAMs engulf developing oligodendrocytes^7^. Given the high purity of the microglial cultures, it is unlikely that contaminating non-microglial cells caused the enrichment of these genes in the WT samples. Their decreased levels in JKO microglia likely reflect the phagocytosis deficit we observed earlier in vitro and in vivo. Gene Ontology (GO) analysis confirmed a reduction in terms associated with neurogenesis and neuronal development (Figure 7D). These and related terms were reported to have increased in microglia following ingestion of apoptotic cells in an in vitro engulfment assay^19^. These findings suggest that Jedi-mediated phagocytosis interacts with and influences neurogenic cells in the neonatal V-SVZ.

**Figure 7.**
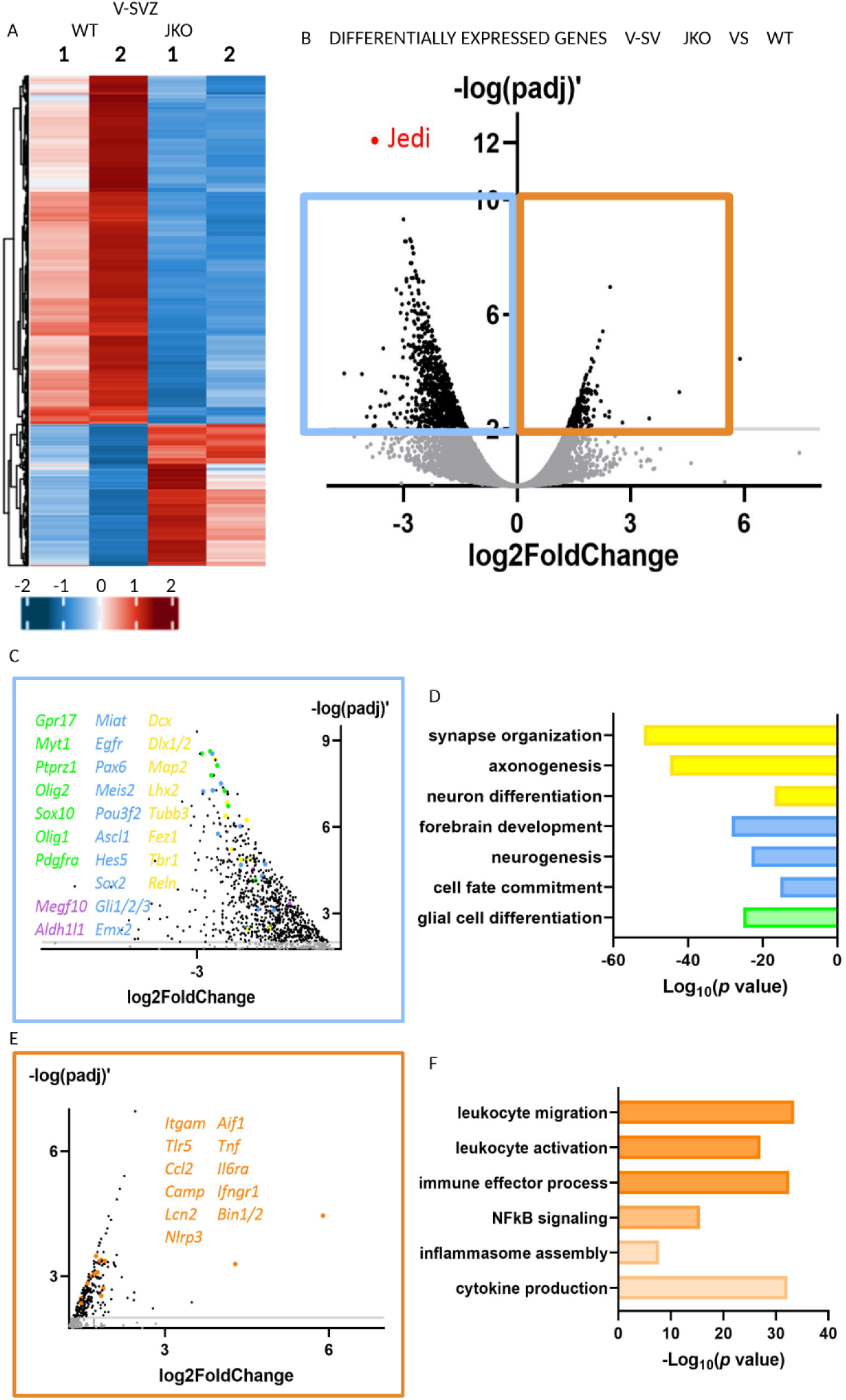
RNA sequencing confirms defective phagocytosis and reveals a neuroinflammatory transformation of Jedi-deficient microglia. 7A. Heatmap demonstrating clear differences in the genetic signature of wild-type (WT) and Jedi knockout (JKO) ventricular-subventricular (V-SVZ) microglia. Colors represent direction of change (blue: relative downregulation, red: relative upregulation) and degree of change (z-scores). 7B. Volcano plot illustrating genes that are downregulated (blue box) and upregulated (orange box) in JKO V-SVZ microglia compared to WT V-SVZ microglia. Threshold of significance: −log(padj)’ ≥ 2.0. 7C. Volcano plot of genes that are downregulated in JKO vs WT (from blue box in 7B). Oligodendrocyte lineage genes (green). Astrocyte genes (purple). Neural precursor cell (NPC) genes (blue). Neuronal development genes (yellow). 7D. Selected downregulated pathways revealed by gene ontology (GO) analysis. Glia-related pathways (green). Neurogenesis-related pathways (blue). Neuronal development pathways (yellow). 7E. Volcano plot of inflammation-related genes that are upregulated in JKO vs WT (from orange box in 7B). 7F. Selected upregulated pathways revealed by GO analysis are highly involved in inflammatory and immune processes.

Of note, we did not find increased expression of genes that had been identified in a previously published single-cell RNA sequencing database of CD11b-expressing sorted microglia as originating from contaminating monocytes/macrophages, neutrophils, or NK cells^7^. However, given the degree of overlap of surface markers between microglia and monocyte/macrophages, we cannot rule out the possibility that some portion of the IBA1^+^ or CD11b^+^ cells that were analyzed via immunofluorescence or RNA sequencing, respectively, are peripheral monocytes/macrophages that have entered the brain.

### Transcriptomic analysis of neonatal Jedi-deficient microglia reveals a neuroinflammatory phenotypic transformation resembling that of neurodegenerative and age microglia

In JKO microglia, we observed increased expression of inflammatory genes such as Camp, Lcn2, Itgam, Tlr5, Ccl2, Aif1, Il6ra, Nlrp3, Ifngr1, Bin1/2, and Tnf (Figure 7E). We did not detect any changes in canonical anti-inflammatory genes (e.g., Il4, Il10, Tgfb1, etc.). GO terms associated with immune cell activation and migration, cytokine synthesis and signaling, and immune responses were enriched in JKO microglia relative to WT microglia (Figure 7F). These data support the idea that Jedi-mediated microglial phagocytosis of apoptotic cells suppresses inflammatory pathways without necessarily promoting anti-inflammatory genes. The changes we observed following Jedi deletion suggest that the microglial population of the neonatal V-SVZ relies on Jedi to help fine-tune inflammatory gene expression and curate its unique genetic signature.

In both the diseased adult brain and aging brain, DAMs and DIMs are present and represent significant deviations from the adult homeostatic microglial phenotype^4^. Loss of the engulfment receptor TREM2 in adult mouse models of Alzheimer’s Disease (AD) and aging mice leads to an expansion of the DIM population at the expense of DAMs, emphasizing the idea that phagocytosis protects against run-away inflammation. We speculated that loss of Jedi would result in a DIM-like phenotype among microglia in the neonatal V-SVZ.

DIMs and DAMs share many of the PAM/DAM markers mentioned above^6^. However, DIMs and DAMs differ in that the DAM developmental program includes substantial downregulation of adult homeostatic genes while this occurs much less robustly in DIMs. We reasoned that we would see little to no differential expression of the PAM/DAM markers between WT and JKO microglia, and we found this to indeed be the case. The adult homeostatic markers listed previously as well as C1qa, Csf1r, Ctss, Entpd1, Fcrls, Hexb, Olfml3, Gpr34, and Sall1 were all significantly enriched in JKO microglia relative to WT microglia (Figure 8A, pink text and data points). In adult microglia, the upregulation of homeostatic markers has been attributed to TGFβ1 signaling^37^, and while we did not see an increase in Tgfb1 transcripts in JKO microglia, Tgfbr1 was elevated relative to WT (log_2_ fold change: 1.52, padj = 0.0023). This suggests that TGFβR signaling may increase following chronic inflammatory pathway activation and constitute an attempt at negative feedback, an idea that has been raised by others in the contexts of DIMs in vivo and of chronic stimulation of microglia in vitro with pro-inflammatory agents^6,38^.

**Figure 8.**
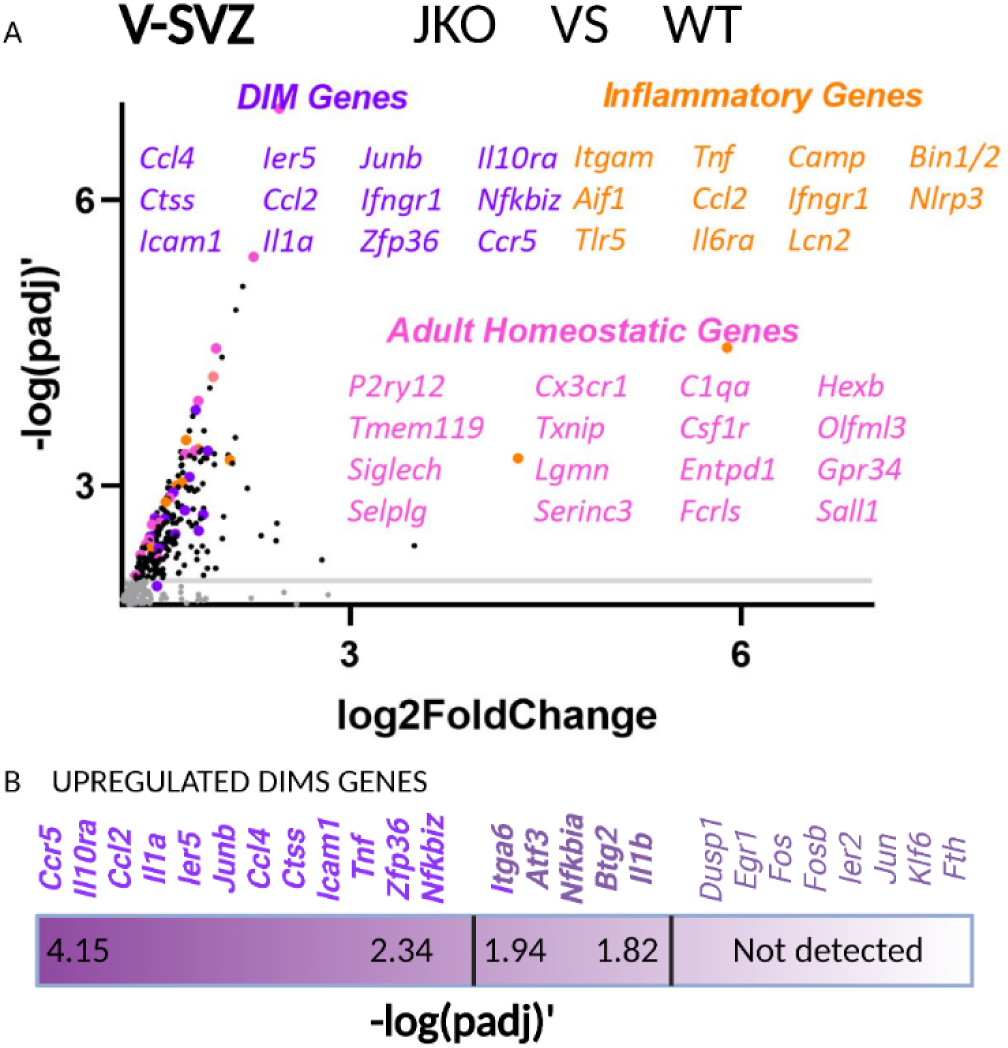
The Jedi-null microglial transcriptome resembles that of inflammatory disease-associated microglia. 8A. Volcano plot of genes that are upregulated in microglia in the ventricular-subventricular zone (V-SVZ) of Jedi knockout (JKO) mice relative to wild-type (WT) mice. Classical inflammatory genes (orange, also shown in Figure 7E), adult homeostatic microglia genes (pink), and disease-associated inflammatory microglia (DIMs) genes (purple). 8B. Heatmap showing 12 of the 25 core DIM markers are significantly upregulated in JKO V-SVZ microglia compared to WT V-SVZ microglia (-log(padj)’ ≥ 2, bolded bright purple, left end of heatmap), 5 of the core markers were increased but did not reach significance (-log(padj)’< 2, bolded dusty purple, middle of the heatmap), and the remaining markers were not detected (light purple, right end of the heatmap). The intensity of the color purple that was used for the heatmap and the text correlates with size of the fold change (dark: higher fold change, light: lower fold change).

The Jedi-deficient microglial genetic and GO signatures resembled those of neurodegenerative microglia (DIMs), which were characterized in an aged AD mouse model and recapitulated in postmortem brain tissue from human AD patients and in aging non-AD brains^5,6^. DIMs are characterized by the expression of inflammation-related genes (e.g., Ccl4, Il1a, Ifngr1, Tnf, etc.), and we found that JKO microglia had a statistically significant enrichment in 12 of the 25 core DIM genes (-log(padj)’ > 2.0, Ccr5, Il10ra, Ccl2, Il1a, Ier5, Junb, Ccl4, Ctss, Icam1, Tnf, Zfp36, and Nfkbiz) (Figure 8B, bright purple bolded text). Additionally, there were compelling upward trends in another 5 of the core DIM genes (1.94 > −log(padj)’ > 1.82; gene name and log2fold change: Itga6 (1.29), Atf3 (1.52), Nfkbia (1.56), Btg2 (1.24), and Il1b (1.59)) (Figure 8B, dusty lavender bolded text). Eight of the 25 core DIM genes were not identified by differential gene expression analysis (Figure 8B, light purple non-bolded text).

We found that many of the GO terms that are enriched in DIMs^6^ were also elevated in JKO microglia (i.e., leukocyte migration & adhesion, IL-6 production, B cell signaling, toll-like receptor signaling, radical oxygen species metabolism, IL-8 production, IL-10 production, nitric oxide biosynthesis, IL-17 production, and acute-phase response). DIMs have higher homeostatic gene expression relative to DAMs and paradoxically upregulate anti-inflammatory genes such as NFκB inhibitors and those involved in IL-10 signaling^6^. In JKO V-SVZ microglia, we observed not only higher levels of adult homeostatic microglial genes (relative to the very low levels seen in PAMs) but also increased expression of NFκB inhibitors and the IL-10 receptor. Taken together, these results argue that V-SVZ-resident microglia in the neonatal brain of Jedi-deficient mice can be characterized as DIMs, which are typically seen during neurodegeneration and in the aging brain.

It has been suggested that DIMs are circulating monocytes that accumulate in the brain over time; an idea that was determined by differential expression of monocyte/macrophage markers (e.g., Cd83, Cd45, Ccr2, etc.)^6,39,40^. In our model, we found no differential expression of transcripts encoding monocyte/macrophage markers between WT and JKO V-SVZ microglia nor between V-SVZ and cortical microglia within the WT group. This suggests that the gene expression profiles in our dataset correspond to microglia rather than to contaminating circulating monocytes/monocyte-derived macrophages.

These data highlight striking similarities between Jedi-deficient neonatal V-SVZ-resident microglia and DIMs, the development of which in the adult brain has already been associated with the loss of the engulfment receptor TREM2. Interestingly, both Jedi and TREM2 signal through Syk recruitment^25,26^. This suggests that Jedi, like TREM2, could play a significant role in determining the overall phenotype of a microglial cell and help to align that phenotype with the surrounding environment and events, particularly as it relates to inflammatory states.

### Cytokine and lipidomic analyses reveal increased IL-1β and lipid mediators in the JKO V-SVZ

Given the neuroinflammatory genetic signature of JKO microglia, we analyzed the levels of several key cytokines via multiplex ELISA in acutely isolated V-SVZ from P7 JKO and WT animals. While IL-6, TNFα, and IL-10 were present in both groups but with no significant differences between genotypes (data not shown), IL-1β levels were significantly elevated in JKO V-SVZ relative to WT V-SVZ (Figure 9A). This is in line with our observations that Nlrp3 of the NLRP3 inflammasome, which cleaves pro-IL-1β into mature IL-1β, is increased in JKO V-SVZ microglia, and that the prevalence of GO terms related to cytokine synthesis and release were increased in the JKO group.

**Figure 9.**
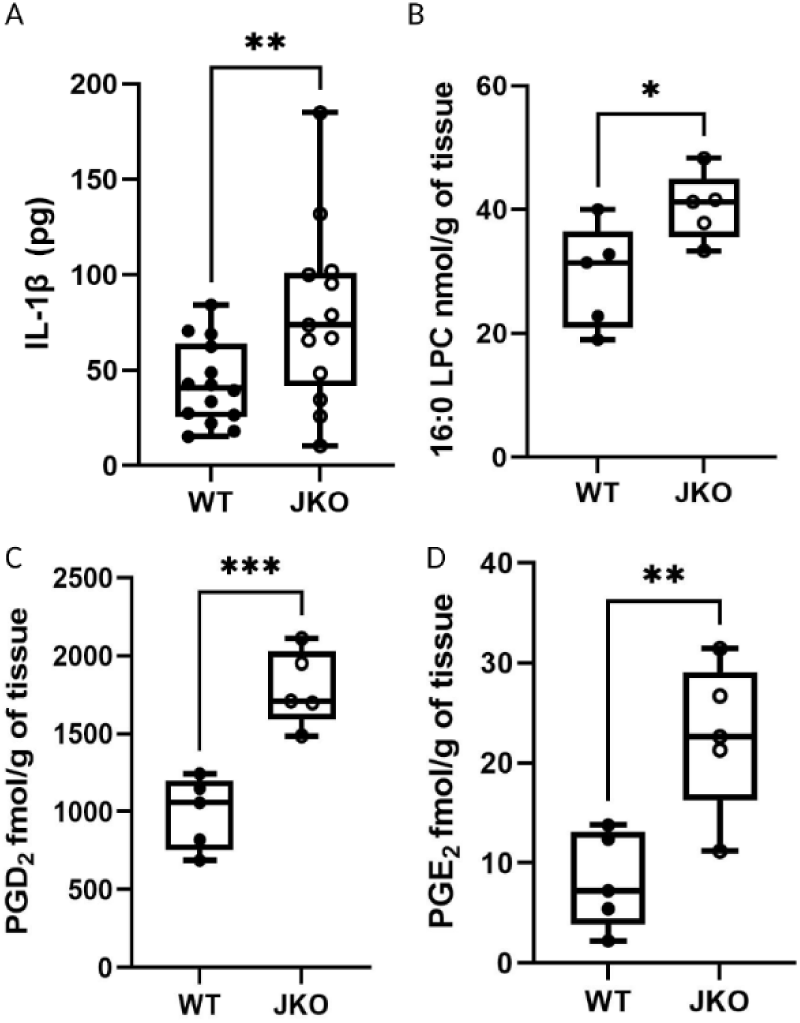
Cytokine and lipidomic analyses reveal increased IL-1β and inflammatory lipid mediators in the JKO V-SVZ. 9A. Picograms of IL-1β Jedi knockout (JKO) and wild-type (WT) ventricular-subventricular zone (V-SVZ). Unpaired one-tailed t test, ** p=0.0025. 9B. Quantification of 16:0 lysophosphatidylcholine (LPC) in JKO and WT V-SVZ. Unpaired two-tailed t test, * p=0.0121. 9C. Quantification of prostaglandin D¬2 (PGD2) in JKO and WT V-SVZ. Unpaired two-tailed t test, *** p=0.0007. 9D. Quantification of prostaglandin E¬2 (PGE2) in JKO and WT V-SVZ. Unpaired two-tailed t test, ** p=0.0069.

In our RNA sequencing dataset, we observed increases in several genes, such as Ptgs1 (cyclo-oxygenase 1, COX-1) and Pla2 family members (Pla2g15, Pla2g3, Pla2g7), that are associated with prostaglandins (PGs) and the metabolism of fatty acids like lysophosphatidylcholine (LPC). To explore whether LPC and PG levels were affected by the loss of Jedi, we performed LC-MS analysis on acutely isolated V-SVZ tissue from P7 WT and JKO mice. The levels of 16:0 LPC, PGD_2_, and PGE_2_ were significantly elevated in JKO V-SVZ relative to WT tissue (Figure 9B–9D). PGF_2_α was detectable, but its levels were not different between genotypes (effect size ±SEM: 5.046 ± 2.740; 95% CI: −1.433 to 11.52; p= 0.1081). An increase in PGs has been associated with inflammation, although the nature of this association may depend on the context^41^. Finally, these changes are likely specific to microglia and not a general phenotype of the JKO mouse because lipidomic analysis of JKO and WT dorsal root ganglia showed no significant differences between groups (A.J. Trevisan and B.D.Carter unpublished data).

Because endocannabinoids have also been implicated in regulating inflammatory pathways in microglia^42^, we measured the levels of several endocannabinoids/endocannabinoid-related lipids (2-arachidonoylglycerol (2-AG), oleoylglycerol (OG), and oleoylethanolamide (OEA)) as well as their precursor, arachidonic acid (AA). We found no significant differences in the concentration of AA, 2-AG, or OG between the two genotypes (data not shown), but OEA was significantly elevated in JKO V-SVZ relative to WT tissue (effect size ±SEM: 4.160 ± 1.605; 95% CI: 0.4585 to 7.862; p= 0.032). OEA is an analog of anandamide, a traditional endocannabinoid, and activates peroxisome proliferator-activated receptor α (PPARα) rather than cannabinoid receptors, the former of which is associated with reducing inflammatory markers in microglia^43^. This elevation in OEA, and thus, the increased possibility of PPARα signaling, could be part of a negative feedback loop that follows chronic inflammation, as was suggested earlier for TGFβ. Taken together, these results further support a phenotypic shift in JKO microglia, one that relates to inflammatory responses.

### Global loss of Jedi reduces V-SVZ neurogenesis

Because JKO microglia resemble DIMs and because inflammation negatively impacts neurogenesis^44–46^, we asked whether NPC proliferation in the V-SVZ was reduced in Jedi-null mice relative to WT mice. We administered a 1-hour EdU pulse in P7 JKO and WT mice and quantified the number of EdU^+^ NPCs, which were identified by fluorescent immunolabeling of the transcription factor MASH1 (ASCL1). MASH1^+^ cells, EdU^+^ cells, and double-labeled cells were present in the V-SVZ of both WT and JKO mice, but all were reduced in JKO relative to WT (Figure 10A–10B, white arrowheads in 10a and 10b indicate EdU^+^MASH1^+^ cells). The reduction in S-phase neurogenic cells in Jedi-deficient animals suggests that microglial phagocytosis plays a role in the typical neurogenic program of the neonatal V-SVZ.

**Figure 10.**
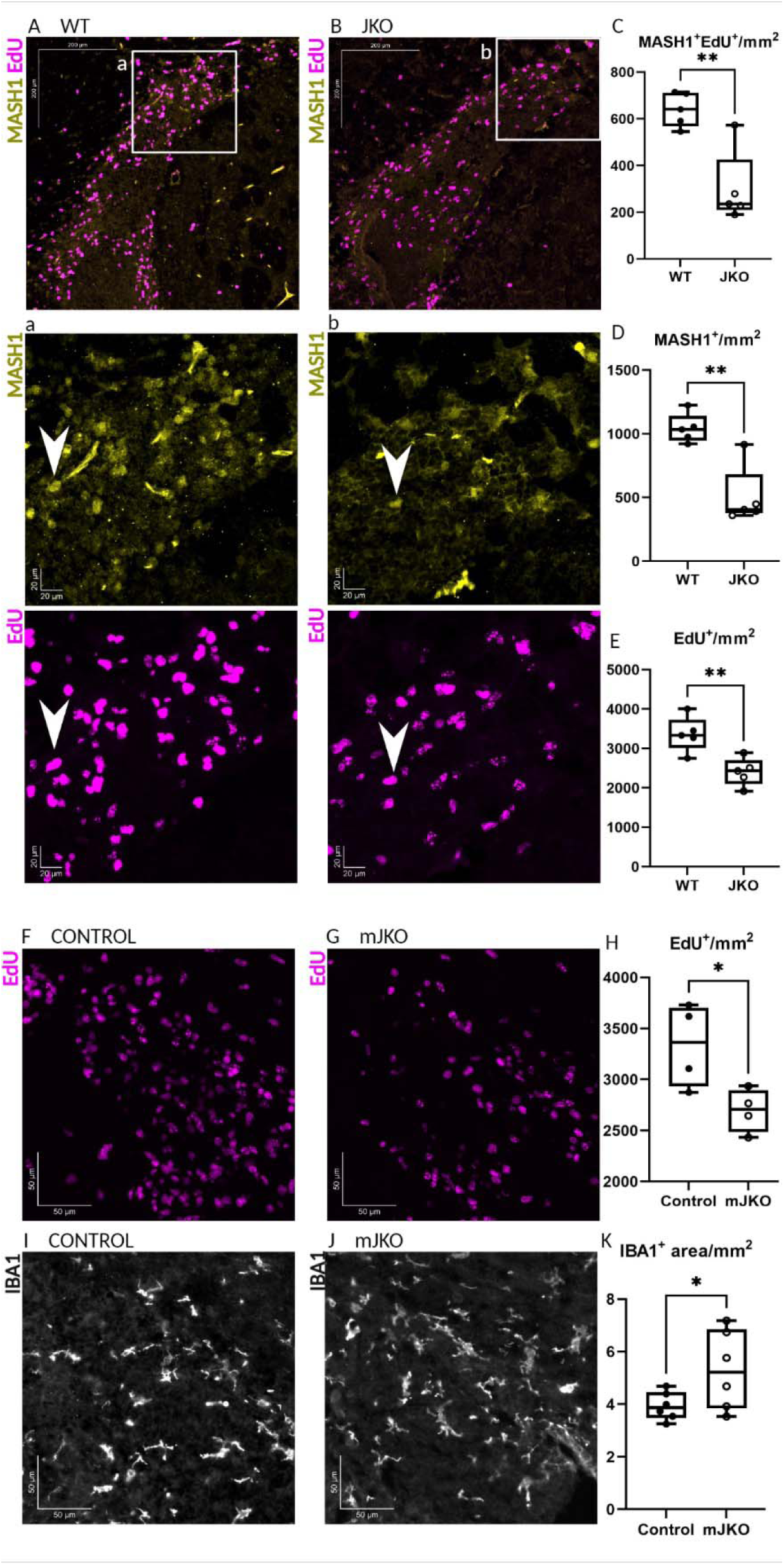
NPC proliferation and differentiation is reduced in the Jedi-deficient V-SVZ, and microglia-specific deletion of Jedi recreates the neurogenic deficit. 10A and 10a. Neural precursor cells (NPCs)(MASH1, yellow) and newly divided cells (EdU (5-ethynyl-2’-deoxyuridine), magenta) in the wild-type (WT) ventricular-subventricular zone (V-SVZ) at postnatal day 7 (P7). The boxed area labeled ‘a’ is displayed in the image immediately below ‘A’, showing MASH1 and EdU separately. White arrowhead indicates an example of a MASH1+EdU+ cells. Scale bar: A, 200 microns (um); a, 20um. 10B and 10b. MASH1+ NPCs (yellow) and newly divided cells (EdU (5-ethynyl-2’-deoxyuridine), magenta) in the Jedi knockout (JKO) V-SVZ at P7. The boxed area labeled ‘b’ is displayed in the image immediately below ‘B’, showing MASH1 and EdU separately. White arrowhead indicates an example of a MASH1+EdU+ cells. Scale bar: B, 200 microns (um); b, 20um. 10C. Quantification of MASH1+EdU+ cells per unit area in the V-SVZ. Unpaired two-tailed t test, ** p=0.0023. 10D. Quantification of MASH1+ cells per unit area in the V-SVZ. Unpaired two-tailed t test, ** p=0.0016. 10E. Quantification of EdU+ cells per unit area in the V-SVZ. Unpaired two-tailed t test, ** p=0.0056. 10F. Newly divided cells (EdU, magenta) in Tamoxifen-injected control animals (Cx3cr1+/+, Cx3cr1CreERT2/+;Jedifl/+, or Cx3cr1CreERT2/+;Jedi+/+) at P7. Scale bar: 50um. 10G. Newly divided cells (EdU, magenta) in Tamoxifen-injected Cx3cr1CreERT2/+;Jedifl/fl microglia-specific JKO (mJKO) animals at P7. Scale bar: 50um. 10H. Quantification of EdU+ per unit area in the V-SVZ area following acute deletion of Jedi from microglia. Unpaired one-tailed t test, * p=0.0163. 10I. Microglia (IBA1, white) in the V-SVZ of Tamoxifen-injected control animals (Cx3cr1+/+, Cx3cr1CreERT2/+;Jedifl/+, or Cx3cr1CreERT2/+;Jedi+/+) at P7. Scale bar: 50um. 10J. Microglia (IBA1, white) in the V-SVZ of Tamoxifen-injected Cx3cr1CreERT2/+;Jedifl/flmJKO animals at P7. Scale bar: 50um. 10K. Quantification of IBA1+ area in the V-SVZ following acute deletion of Jedi from microglia. Unpaired one-tailed t test, * p=0.0306.defect in Jedi-null mice.

### Microglia-specific loss of Jedi is sufficient to reduce V-SVZ neurogenesis

Because Jedi is also expressed in blood vessels (Figure 1), we wanted to determine whether acute loss of Jedi specifically in microglia would be sufficient to reduce neurogenesis. We used Cx3cr1^CreERT^2^/+^ mice crossed to Jedi^fl/fl^ mice to generate a Tamoxifen-inducible, microglia-specific Jedi KO line (mJKO). Recombination was activated in pups by injecting the dams twice daily with Tamoxifen starting 1 day after birth for a total of 3 days. Using qPCR in immunomagnetically sorted microglia from the micro-dissected V-SVZ, we found this approach was sufficient to significantly decrease Jedi mRNA (Supplementary Figure 4). At P7, we evaluated proliferation in the V-SVZ using EdU. In agreement with the global JKO, we found that loss of Jedi in microglia alone was sufficient to reduce the number of EdU^+^ cells in the V-SVZ (Figure 10F–10H). Additionally, we observed an increase in IBA1^+^ area (Figure 10I–10K), which we previously found was correlated with inflammatory microglial phenotype (Figures 4 and 5). These results indicate that Jedi cell-autonomously regulates microglial phagocytosis in a way that indirectly supports neurogenesis.

Given that IL-1β was significantly elevated in the JKO V-SVZ and is detrimental to neurogenesis, causing cell cycle arrest and apoptosis via direct IL-1 receptor signaling in NPCs^47^, we sought to rescue the neurogenesis defect via intracerebral (i.c.) injection of the recombinant human IL-1 receptor antagonist (IL-1RA). At P4, WT and JKO pups received a single i.c. injection of IL-1RA in the right hemisphere, and on P7, the pups were euthanized following a 1-hour EdU pulse (Supplementary Figure 5A–C). It has been shown that substantial suppression of cytokine signaling in the perinatal V-SVZ causes excessive NPC proliferation^15^, and over-labeling of the proliferating cells could interfere with accurate quantification, so the length of the EdU labeling reaction time was reduced from thirty minutes to fifteen minutes. Decreasing the EdU labeling time successfully halved the number of labeled EdU^+^ cells (Supplementary Figure 5D). Quantification of EdU-labeled cells in the V-SVZ of WT and JKO mice revealed that IL-1RA injection abrogated the previously observed reduction of proliferating cells in JKO such that there was no longer any significant difference between WT and JKO (Figure 11A–11C). These data support the idea that reduced neurogenesis in Jedi-null animals is driven by an increase in IL-1β signaling, and that increased IL-1β signaling is part of a neuroinflammatory phenotypic transformation that occurs when the engulfment of apoptotic neurogenic cells is disrupted.

**Figure 11.**
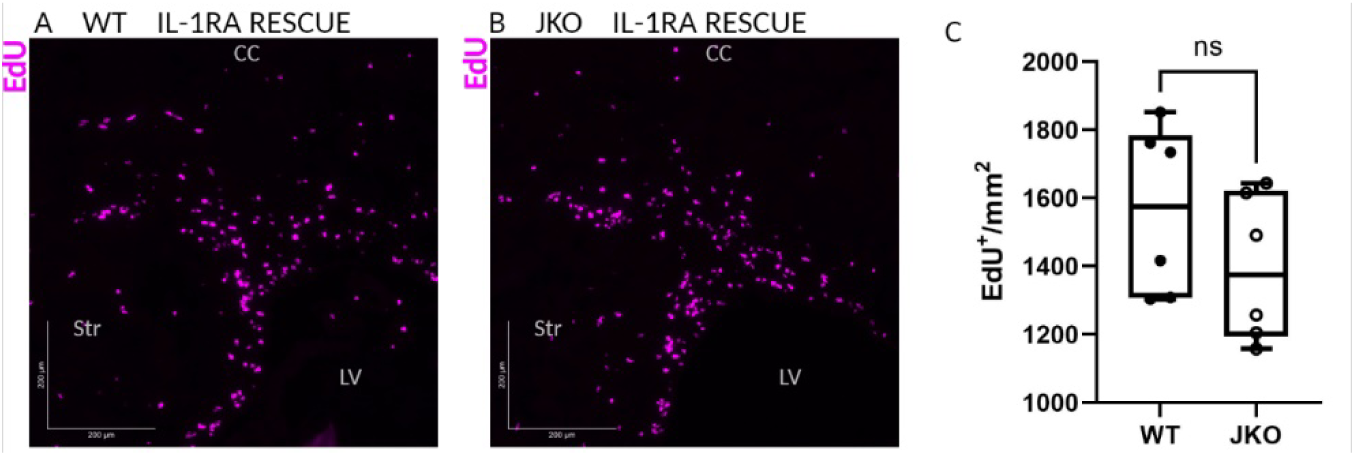
Inhibition of IL-1R signaling rescues the neurogenesis defect in Jedi-null mice. 11A. Newly divided cells (EdU (5-ethynyl-2’-deoxyuridine), magenta) in the wild-type (WT) ventricular-subventricular zone (V-SVZ) at postnatal day 7 (P7) following interleukin-1 receptor antagonist (IL-1RA) intracerebral (i.c.) injection at P4. Corpus callosum (CC), Striatum (Str), and Lateral Ventricle (LV) are indicated for anatomical reference. Scale bar: 200 microns (um). 11B. Newly divided cells (EdU, magenta) in the Jedi knockout (JKO) V-SVZ at P7 following interleukin-1 receptor antagonist (IL-1RA) i.c. injection at P4. Corpus callosum (CC), Striatum (Str), and Lateral Ventricle (LV) are indicated for anatomical reference. Scale bar: 200 microns (um). 11C. Quantification of EdU+ cells per V-SVZ area following IL-1RA rescue. Unpaired one-tailed t test, not significant, p=0.1193.

## Discussion

Microglia are key players in regulating homeostasis in the healthy CNS as well as in pathological conditions. These cells are exquisitely sensitive to their environment, in part through the material they phagocytose, and they rapidly modify their phenotype in order to address local changes. Here, we identified the phagocytic receptor Jedi as a critical regulator of microglial phenotype in the V-SVZ, and a crucial indirect modulator of neurogenesis. Specifically, we found Jedi expressed by microglia in the V-SVZ neurogenic niche at P7, which coincides with the presence of microglial with a specialized phenotype. We demonstrate that Jedi mediates microglial phagocytosis, and Jedi deletion disrupts not only phagocytosis but also the properties of these unique microglia, leading to morphological, functional, and transcriptomic changes that are consistent with those seen in microglia in the diseased and aging brain. In the Jedi-null microglia, there was an increase in cytokine synthesis and signaling pathways, most notably, elevated levels of IL-1β, which we demonstrate suppresses neurogenesis. Overall, our findings illustrate that Jedi, a novel engulfment receptor in the CNS, is at the core of how microglia interact with the neurogenic environment and fine-tune their behavior to support neurogenesis.

### Phagocytic receptor expression and function in V-SVZ microglia is a matter of debate

Only a few studies have explicitly examined the influence of microglial phagocytosis on neurogenesis. Most in vivo work has focused on the deletion of the TAM receptors (Tyro3/Axl/MerTK) and to some extent, P2RY12. All reached slightly different conclusions, likely due to methodological differences as well as to biologically relevant differences between the ages and brain regions examined in each study. Consequently, whether or how microglial phagocytosis impacts neurogenesis in the V-SVZ is unclear. Fourgeaud et al. reported that adult (3-6-months-old) Axl/Mertk-null mice accumulated dead cells in the V-SVZ and rostral migratory stream (RMS) but had more newly generated neurons in the olfactory bulb^18^. This raised the possibility that the selective and active culling of migrating neuroblasts, a process termed ‘phagoptosis’, is a component of microglia function in the RMS and that loss of microglial phagocytosis resulted in a net increase in neuroblast survival. Regardless of whether microglial phagoptosis is or is not part of normal V-SVZ neurogenesis, it does not explain the increase in apoptotic cells in the V-SVZ observed in TAM KO mice. Whether Axl/Mertk deletion affected NPC proliferation, differentiation, or survival specifically in the V-SVZ was not addressed.

Ribeiro Xavier et al. reported that microglia in the V-SVZ and RMS of juvenile mice (1-2-months-old) showed no evidence of phagocytosis of neurogenic cells, and in fact, TREM2 and P2Y-family receptor expression in microglia was absent or very low in these regions^16^. TAM receptor expression was not examined. When microglia were depleted from the V-SVZ, there was an increase in BrdU^+^ cells in the V-SVZ and RMS, echoing the observations made by Fourgeaud et al. in the olfactory bulb. Crucially, however, a significant portion of the BrdU^+^ cells had pyknotic nuclei, signaling cell death. These findings were interpreted as reflecting disrupted neuroblast migration and increased cell death, independent of phagocytosis, which was not detected in the normal V-SVZ and RMS. Counterintuitively, this failure to detect phagocytosis may have resulted from the efficiency and speed of dead cell clearance, which was demonstrated by Sierra et al. to start and end in as little as thirty minutes in the healthy wild-type adult SGZ^48^. This raises the possibility that the increase in pyknotic newborn cells reflects reduced clearance of dead cells rather than or in addition to reduced survival. We encountered this same limitation in our work, though the targeted deletion of an engulfment receptor rather than the removal of all V-SVZ microglia favors the ‘reduced clearance’ explanation. Studies aimed at addressing this confusion between increased death and reduced clearance are underway.

Microglial expression of phagocytic receptors and the fact that its disruption affects neurogenesis are less debated in the SGZ, though the nature of microglial influence on neurogenesis is still not clear. Diaz-Aparicio et al. demonstrated that deletion of Mertk/Axl or P2ry12 reduced the number of neuroblasts generated in the adult SGZ^19^. However, P2ry12 KO mice also displayed a reduction in the number of proliferative NPCs/neuroblasts, while Mertk/Axl KO mice did not, and Mertk/Axl KO mice displayed increased apoptosis in the SGZ without affecting proliferation, which was not the case for P2ry12 KO mice. The reason for these differential outcomes of phagocytic receptor knockout is not yet understood but likely reflects the idea that engulfment receptors are not redundant and each serves a specific purpose at a specific time and place. Unfortunately, very little of this idea has been explored. Ji et al. also examined proliferation and survival of NPCs in the adult SGZ of TAM receptor KO animals and found that loss of TAM receptors reduced NPC proliferation and survival, and, independently, reduced neuronal differentiation^49,50^. Despite their differences, these studies provided support for the linkage of phagocytosis to NPC proliferation, differentiation, and survival, even if the mechanisms behind these links were not fully identified.

### Microglial influence on neurogenesis relies on phagocytosis-dependent changes to the ‘secretome’

Several studies point to the body of secreted factors as a key determinant of the phenotypic changes that occur after phagocytosis. Fortunately, significant progress has been made in characterizing the composition of the phagocytic secretome, which includes cytokines, signaling lipids, growth factors, and more^19^.

Cytokines are the most likely mechanism of microglial influence on NCP function, and IL-1β has been implicated in microglia-associated changes in neurogenesis. For example, TAM receptor-null microglia increased the expression of canonical inflammatory molecules, including IL-1β, which coincided with decreased SGZ neurogenesis^49^, and persistent overexpression of IL-1β in WT adults impaired hippocampal neurogenesis^51^. In vitro, microglia-derived IL-1β promoted neurogenesis at low concentrations (1ng/mL) but had the opposite effect at higher concentrations (10ng/mL)^38,44^. In agreement with this, we showed that IL-1β levels were elevated in the JKO V-SVZ, which correlated with reduced neurogenesis, and blocking IL-1β signaling rescued NPC proliferation in JKO mice. Our experimental design permitted us to trace the reduction in neurogenesis and concomitant increase in IL-1β and IL-1R signaling back to disrupted phagocytosis that resulted from the loss of Jedi specifically in microglia.

The idea that disrupted phagocytosis would lead to pathological levels of IL-1β is supported by several studies. In cultured microglia exposed to LPS, the typical robust expression of IL-1β, IL-6, and TNFα was significantly dampened by pre-treatment of the microglia with dead cell debris, and this suppression of inflammatory cytokines occurred without necessarily an accompanying increase in TGFβ or IL-10 production, though this may be context-specific: increases in TGFβ, PGE_2_, and nerve growth factor (NGF) have also been observed after LPS stimulation of microglia that have ingested apoptotic cells^52–55^. Our observations agree with these findings, emphasizing the role that Jedi plays in transducing an apoptotic cell-derived signal into an immunomodulatory phenotype that supports neurogenesis by mitigating the potentially neuroinflammatory effects of high levels of dead cells.

Although blocking IL-1β rescued the deficit in NPC proliferation in JKO mice and pointed to the suppression of the IL-1 axis as a mechanism for supporting the neurogenic cascade, we cannot rule out the possibility that other factors released by microglia also play a role. Additional mechanisms may involve lipid mediators, which we found to be altered in Jedi-deficient mice. Our lipidomic analysis focused on prostaglandins, which are members of the eicosanoid family of arachidonic acid-derived lipid mediators, including thromboxanes and prostacyclins, all of which influence inflammation, sometimes in opposing ways^56^. Concomitant with the increase in IL-1β levels and the emergence of a neuroinflammatory genetic signature in JKO microglia, we saw a significant elevation in PGD_2_ and PGE_2_.

In macrophages, ingestion of apoptotic cells induces COX-2-dependent production of PGE_2_, which in turn promotes the secretion of TGFβ, inducing a ‘tolerogenic’ phenotype^57^. We were unable to reliably detect TGFβ in our V-SVZ lysates and our sequencing dataset did not show differential expression of Tgfb or Ptgs2 (COX-2) in JKO microglia. On the other hand, microglia-derived PGD_2_ has been implicated in inflammation, demyelination, and astrogliosis both in vitro and in vivo^58^. PGD/E_2_ can also be synthesized by COX-1 activity, and COX-1-derived PGs may have functions different from those of COX-2-derived PGs^59^. We found that Ptgs1 was indeed upregulated in JKO microglia. Whether these lipid mediators contribute to the neuroinflammatory phenotype and subsequent neurogenesis defect or whether they play a part in a negative feedback loop (like that proposed to explain the increase in OEA, Tgfbr1, and Il10ra) is not yet known. We also found an increase in LPC, which is notable because LPC is associated with NLRP3 inflammasome-dependent IL-1β synthesis^60,61^. We limited our analysis to several key lipid species, but given our current findings, a more comprehensive lipidomic analysis in the future will be beneficial to understanding the role that lipid mediators play in the Jedi-deficient microglial phenotype and neurogenesis deficit.

Changes in trophic factor release may also be relevant to how microglial phagocytosis influences neurogenesis. An in vitro engulfment assay using dead cells as bait found that of the 224 genes that were differentially expressed in phagocytic microglia and had already been identified as regulators of neurogenesis, the most abundant genes were those of growth factors and cytokines (51 and 39 genes, respectively), though the largest fold-changes were among peptides and hormones^19^. In our dataset, we found that only 12 of those 224 genes were differentially expressed between JKO and WT V-SVZ microglia. Seven of those were chemokines and cytokines that were found to be downregulated after phagocytosis induction in vitro, and consistent with this observation, were significantly upregulated in our dataset of JKO V-SVZ microglia. Thus, our results do not support changes in growth factor expression as major contributors to the microglial phenotype or neurogenesis deficit in Jedi-null mice, although it is possible that there are alterations in the release of such factors or the distribution and expression of their receptors.

### Jedi provides a long-sought mechanistic connection between phagocytosis and neurogenesis

Jedi has not been extensively studied in the nervous system. In fact, its role has only been characterized in the peripheral nervous system, in dorsal root ganglia^24^. According to single-cell RNA sequencing datasets, PAMs co-express a variety of engulfment receptors^5^, though for most, their expression profile and functions have not been explored. Our GO analysis of JKO microglia returned six GO terms involving phagocytosis (whether the fold change was positive or negative was gene-specific); however, there was no significant functional compensation for the absence of Jedi. In both our in vitro and in vivo engulfment assays, we observed the presence of phagocytic microglia in the JKO samples, which is consistent with previous findings that deletion of Axl/Mertk or P2ry12 did not completely abrogate microglial phagocytic ability. This supports the idea that microglia have many tools for clearing apoptotic or extracellular debris, and it highlights how important clearance is for brain homeostasis. The various engulfment receptors expressed by microglia are likely independent and/or complimentary and may provide additional precision in a microglial cell’s response to specific stimuli. Additionally, engulfment receptors belong to an array of families and activate different signaling modalities that likely further increase the granularity and relevance of microglial interactions with the events occurring around them. For example, Axl and MerTK are receptor tyrosine kinases, P2RY12 is a G-protein-coupled receptor, TREM2 signals via immunoreceptor tyrosine-based activation motifs (ITAMs) in its co-receptor, DAP12, and similarly, Jedi signals via ITAMs in its intracellular domain.

Despite their differences, all of these receptors can activate the soluble tyrosine kinase Syk^25,62–64^, suggesting that Syk may be a common signaling node. How and why the pathways converge or diverge following Syk activation-or if they occur in parallel-is not clear. Nevertheless, Syk promotes cytoskeletal remodeling^65^, which is necessary for several steps in the phagocytic process and would thus be predicted to be a shared signaling pathway. In addition, it was recently reported that Syk was necessary for TREM2-mediated conversion of microglia to the DAM phenotype^26,66^, suggesting that Syk activation alters microglial function much more broadly than just reorganizing the cytoskeleton. Despite their shared involvement of Syk, the receptors likely bind a diverse set of ligands and activate common as well as unique signaling pathways, allowing microglia to mount a response tailored to their local environment. TREM2 is the most striking example of the role played by phagocytosis in generating a context-specific microglial phenotype. It was found that while the DAM phenotype is TREM2-dependent, the PAM phenotype, despite its similarities to that of DAMs, is TREM2-independent^7^, and our findings suggest that it may instead be Jedi-dependent, as so, Jedi may be the first step in an identity-defining process necessary for microglia function in the developing brain.

Beyond more deeply understanding the signaling events that follow Jedi activation in neonatal microglia, the extent to which changes in Jedi expression correlate with the response to or onset and progression of neuropathology in the adult and aging brain should be explored. Pharmacological targeting of engulfment receptors is gaining significant attention in pre-clinical and clinical research^67,68^, and it may prove beneficial, as it has for TREM2^69–71^, to explore Jedi in these spheres.

## Materials and methods

### Experimental design and statistical analysis

Comparisons were made between wild-type (WT) and constitutive Jedi knockout (JKO) mice, or between Cx3cr1^CreERT2/+^;Jedi^flox/flox^ (microglia-specific JKO, mJKO) mice and littermate controls (see genotypes in Mice section). Males and females were equally represented in the analyses. The experimenter was blinded to genotype until all the raw values (e.g., engulfment index, cell number, puncta number, area, etc.) were collected and recorded. The in vitro bead engulfment assay was repeated three times and the data points graphed in Figure 2 are the combined results of those three independent experiments. Each chamber of the 4-well chamber slide was divided into six regions and one image was taken per region. Additional images were taken if, for a given condition, six images did not provide at least fifteen cells for analysis. With respect to immunofluorescent labeling and cytokine analysis, biological replicates originated from different litters, and each experiment contained 4-6 mice per group. Within an experiment, 3-4 slides were included for each animal, each slide containing 2-3 sections. Two images per region analyzed were acquired per section. The values that appear in the graphs are averages of values quantified from all the images for each animal. For the measures of microglial morphology, fifteen individual cells were selected at random in each image. In vivo measures were divided by the area (or in the case of confocal imaging, the volume) of the defined region of interest (ROI) to generate a per mm^2^ or mm^3^ value. Statistical analyses were performed using GraphPad Prism 9, and specific tests are indicated in the figure legends. Exact p values will be included when available; but in their absence, the following notations are used to indicate the statistical significance of each test: *p < 0.05, **p < 0.01, ***p < 0.001.

#### Choice of statistical tests

Both one-and two-tailed t tests are used. One-tailed tests were selected when the hypotheses were directional (i.e., Alternative hypothesis (H_1_): The number of TUNEL^+^ cells is higher in the V-SVZ of JKO vs WT. Null hypothesis (H_0_): The number of TUNEL^+^ cells is not higher in V-SVZ of JKO vs WT.) Two-tailed tests were used when the hypothesis was that the measure would be different between JKO and WT groups but the direction of the change could not be hypothesized (i.e., Alternative hypothesis (H_1_): The number of TUNEL^+^ cells in the cortex and striatum is different in JKO vs WT. Null hypothesis (H_0_): The number of TUNEL^+^ cells in the cortex and striatum is not different in JKO vs WT.) In some cases, the distribution of the data points prompted the use of post-hoc normality tests (i.e., Shapiro-Wilk and Kolmogorov-Smirnov). If the data were not normally distributed, a Mann-Whitney U test was used to compare the two groups in question.

### Mice

All animal procedures were performed in accordance with the Vanderbilt University Medical Center’s comprehensive Animal Care and Use Program (ACUP) and NIH guidelines for the Care and Use of Laboratory

#### Animals

Mice were housed under a controlled 12-hour light/dark cycle and fed standard laboratory rodent diet (LabDiet, Cat #. 5001) with water available ad libitum. Jedi knockout (JKO) mice were Pear1^tm1a(KOMP)Wtsi^ mice derived from embryonic stem cells provided by the International Mouse Phenotype Consortium (IMPC, Cat # CSD31459_C05). Control mice were wild-type (WT) C57/BL6 mice obtained from Jackson Labs and then maintained by our laboratory (catalog no. 000664). Pear1^tm1a(KOMP)Wtsi^ mice were crossed with mice expressing the FLP1 recombinase driven by the Gt(ROSA)26Sor promoter. This cross resulted in deletion of the frt-flanked sequence, thereby producing a wild-type Jedi1 allele with exons 3 and 4 floxed (Jedi1^fl/fl^). Microglia-specific JKO (mJKO) were generated by breeding B6.129P2(Cg)-Cx3cr1^tm2.1(cre/ERT2)Litt^/WganJ (RRID:IMSR_JAX:021160) with Jedi^fl/fl^. The Cx3cr1^CreERT2^ mice are knock-in/knock-out mice expressing Cre-ERT2 fused with EYFP, thus they were bred as heterozygotes to avoid generating Cx3cr1^-/-^ mice. Cx3cr1^CreERT2/+^;Jedi^fl/+^ mice were crossed with Jedi^fl/+^ mice ensuring a litter of mixed genotypes. Littermates with the following genotypes were used as controls: Cx3cr1^+/+^, Cx3cr1^CreERT2/+^;Jedi^fl/+^, or Cx3cr1^CreERT2/+^;Jedi^+/+^.

#### Genotyping

Mice that were used for genotyping were ear-tagged, and ear snips were collected. Ear snips were heated at 93°C for one hour in 25mM NaOH, 0.2mM EDTA, then 40mM Tris-HCl was added at 1:1 to stop the DNA extraction. Samples were stored at −20°C until genotyping. The primers, annealing temperature, extension time, and product sizes are listed in Table 1. The reaction was done using GoTaq® Flexi (Promega, Cat # M829) on a GeneAmp® PCR System 2700 (Applied Biosystems). PCR products were run on a 4% agarose-TAE gel.

**Table 1.**
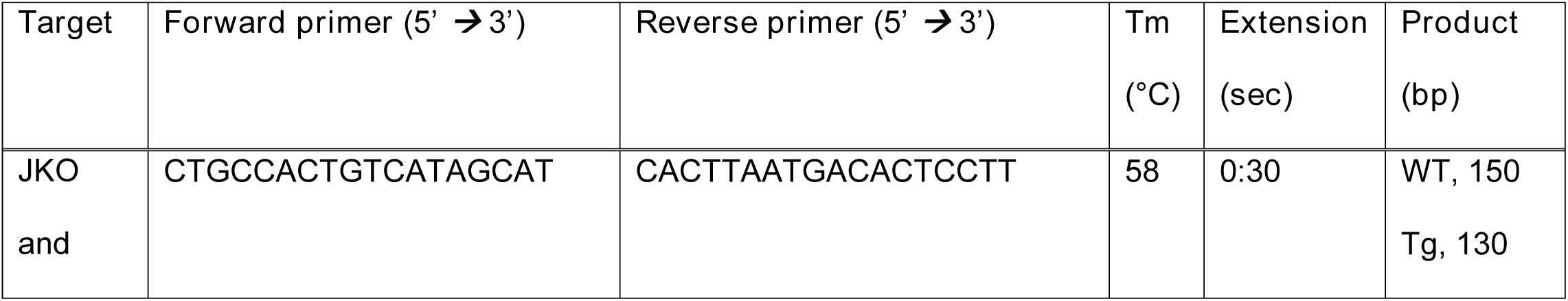
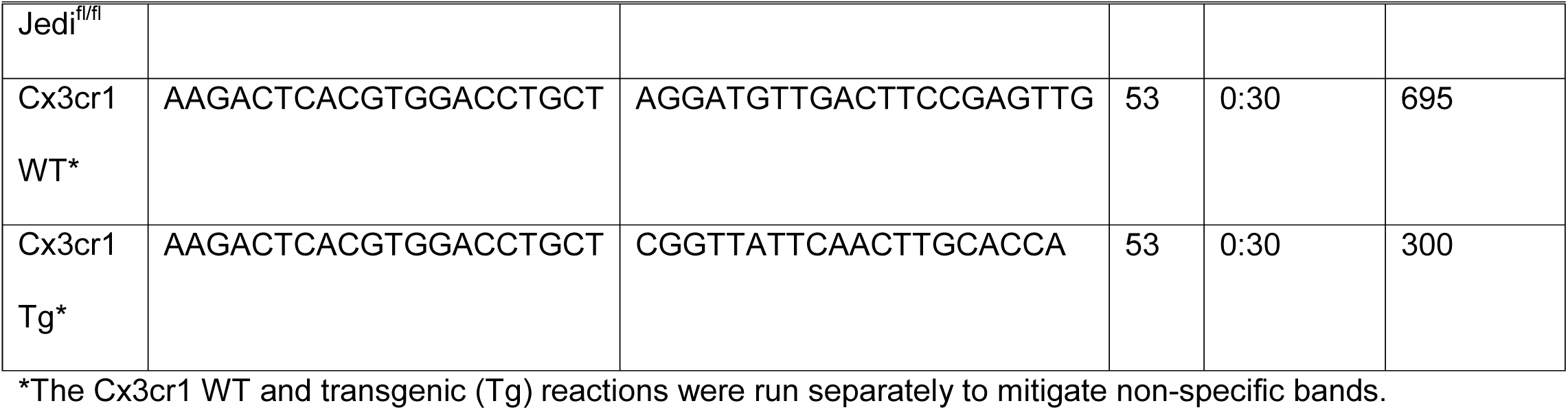
Genotyping primer sequences

### Primary microglia cultures

Postnatal day 7 (P7) pups were anesthetized on ice for 5-10 minutes. After passing the toe pinch test, the animals were decapitated. The brain was removed from the cranium and the cortical hemispheres, excluding the hippocampi, were isolated. The tissue was roughly chopped with a sterile blade and enzymatically dissociated with the Papain Dissociation system, according to the manufacturer’s instructions (Worthington Biochemical Corporation, Cat # PDS). Cortical mixed glia were seeded on 10 microgram/milliliter (ug/mL) poly-D-lysine-coated T25 flasks (minimum two brains per flask) and grown in DMEM/F12+GlutaMAX™ (Gibco, Cat # 10565-018) supplemented with 10% fetal bovine serum and 1% Pen/Strep until confluency (∼1 week). To harvest microglia, the flasks were secured in a secondary container that limits sliding and jostling of the flask during shaking. Shaking took place in a 37°C shaker for five hours at 175 revolutions per minute. The media, now containing microglia, was removed into a new tube and microglia were pelleted down (70 x g for five minutes) and all the media except for 100uL was pipetted off; the cell pellet may not be visible. The cells were resuspended in 1mL of media and 250uL of the cell suspension was placed in each chamber of 4-well chamber slide. The bead engulfment assay was performed the following day.

### In vitro bead engulfment assay and staining

FluoSpheres™ Carboxylate-modified microspheres (Invitrogen, Cat # F8826) were vigorously vortexed for one minute then diluted 1:33 in 0.5% bovine serum albumin (BSA). This stock was then further diluted 1:30 in media (10% FBS, 1% Pen/Strep, DMEM/F12+Glutamax) to yield the bead working stock. The media in the chamber was aspirated and replaced with the bead working stock, taking care to triturate the bead working stock several times between chambers to resuspend any beads that may have settled in the tube. The microglia were incubated at 37°C with the beads for four hours, two hours, and five minutes (0-hour time point), then the bead working stock was pipetted off and the cells were gently rinsed with 1x PBS. Cells were fixed in 10% neutral buffered formalin (NBF) for ten minutes at room temperature, then washed three times, three minutes per wash in 1x PBS. Cells were permeabilized for three minutes at room temperature using 0.2% Triton-X100 in 1x PBS, then washed three times, three minutes per wash in 1x PBS, before undergoing blocking for fifteen minutes at room temperature (blocking solution: 5% BSA, 0.1% Tween-20 in 1x PBS). Phalloidin-488 methanolic stock (Invitrogen, Cat # A12379) was diluted 1:40 in 1x PBS, and the cells were incubated with the phalloidin-488 working solution for sixty minutes at room temperature in the dark. Cells were washed three times, three minutes per wash in 1x PBS. The cells were counterstained with ProLong Gold with DAPI (Invitrogen, Cat # P36931), covered with a glass coverslip, and left at room temperature in the dark until imaging.

### EdU injections

The reconstituted EdU stock solution (5 milligram/milliliter (mg/mL) was diluted in sterile 1x PBS to a working solution of 1.25mg/mL. P7 mice were weighed to calculate the volume necessary to achieve a dose of 15mg/kilogram (mg/kg). Mice were then injected subcutaneously in the skin fold above the neck using a 300uL 29G1/2 syringe (Comfort Point, Cat # 26018); so that each mouse had precisely one-hour exposure to EdU, animals were injected 15 minutes apart to allow time for anesthesia, perfusion, and brain removal after euthanasia.

### Tissue processing

#### Paraffin-embedded sections

The animals were anesthetized on ice and underwent transcardial perfusion with 5mL 1x PBS followed by 5mL 10% NBF. Once extracted, the brains were placed in 10% NBF overnight at room temperature on a rocker. The following day, the brains were washed three times for twenty minutes in 1x PBS, underwent processing for paraffin embedding, and 7-micron-thick sections were collected using a sliding microtome. Slides were stored at room temperature until immunofluorescent labeling.

#### Frozen sections

The animals were anesthetized on ice and underwent transcardial perfusion with 5mL 1x PBS followed by 5mL 4% paraformaldehyde (PFA). Once extracted, the brains were placed in 4% PFA in 1x PBS overnight at room temperature on a rocker. The following day, the brains were cryoprotected in 30% sucrose for 24 hours or until the samples sank to the bottom of the tube. Brains were embedded in optimal cutting temperature medium and 10-micron-thick sections were collected, placed on Fisherbrand™Superfrost™ Plus slides (Fisher Scientific, Cat # 12-550-15) and stored at −80°C. Fifty-micron-thick sections were collected and stored at 4°C in 1x PBS in a 24-well plate sealed with Parafilm®.

### Immunofluorescence

#### Paraffin-embedded sections

Slides were baked at 65°C for thirty minutes. Paraffin was removed by serial 5-minute incubations in xylenes and decreasing concentrations of ethanol. Slides were rinsed for ten minutes in distilled water, then washed in 1x PBS. Antigen retrieval was performed in a pressure cooker for fifteen minutes (manual/default mode, no special program). For IBA1, slides were submerged in citrate buffer pH 6.0 (0.01M anhydrous citric acid, 0.05% Tween-20, in distilled water). For MASH1, Tris-EDTA pH 9.0 was used (0.01M Tris Base, 1.26mM EDTA, 0.05% Tween-20, in distilled water). Slides were rinsed once with 1x PBS, then blocking solution (5% BSA, 0.1% Tween-20, in 1x PBS) was applied to the sections for one hour before proceeding to the primary antibody overnight at 4°C or directly to EdU labeling. Primary antibodies: Polyclonal Rabbit anti-Mouse IBA1 (Wako, Product Number 019-19741; used at 1:750 in 1x PBS); Monoclonal Mouse anti-Rat MASH1 (BD Pharmigen, Cat # 556604; used at 1:200 in blocking solution). Slides were washed twice, three minutes per wash, in wash buffer (0.05% PBS-Tween), then secondary antibody was applied for one hour at room temperature in the dark. Secondary antibodies: Goat anti-Rabbit 488 (Invitrogen, Cat # A-11008; used 1:1000 in 1x PBS), Donkey anti-Mouse 488 (Invitrogen, Cat # A-21202; used at 1:1000 in 1x PBS). Slides were washed twice, three minutes per wash, in wash buffer. Mounting media was applied, a glass coverslips placed on top, and slides were left to dry in the dark at room temperature until imaging.

#### Frozen sections

Ten-micron-thick slide-mounted sections. Slides were post-fixed at room temperature in 4% PFA for thirty minutes then washed once for five minutes in 1x PBS. Sections were permeabilized with 0.2% Triton-X100 for ten minutes at room temperature, then rinsed in 1x PBS before blocking for thirty minutes at room temperature followed by the primary antibody overnight at 4°C. Primary antibodies: Polyclonal Sheep anti-Mouse PEAR1 (R&D Systems, Cat # AF7607; used at 1:50 in blocking solution); Polyclonal Rabbit anti-Mouse IBA1 (Fujifilm Wako Chemicals, Cat # 019-19741; used at 1:500 in blocking solution). Slides were washed twice for three minutes in wash buffer, then received secondary antibodies for one hour at room temperature in the dark. Secondary antibodies: Donkey anti-Sheep 568 (Abcam, Cat # ab175712; used at 1:1000 in blocking solution); Donkey anti-Rabbit 488 (Invitrogen, Cat # A-21206; used at 1:1000 in blocking solution); Goat anti-Rabbit 647 (Invitrogen, Cat # 21245; used at 1:1000 in blocking solution). Slides were washed twice, three minutes per wash, in wash buffer. Mounting media was applied, a glass coverslips placed on top, and slides were left to dry in the dark at room temperature until imaging.

#### Fifty-micron-thick free-floating sections

Sections were post-fixed for one hour at room temperature in 4% PFA, then washed once for five minutes in 1x PBS. Blocking was performed for one hour at room temperature (blocking solution: 5% BSA, 0.3% Triton-X100, in 1x PBS), then the slides received primary antibody for 48-72 hours at 4°C. Primary antibody: Rabbit anti-Mouse IBA1 (same as for 10um-thick sections). Sections were washed twice for three minutes in wash buffer before receiving the secondary antibody for two hours at room temperature in the dark. Secondary antibody: Donkey anti-Rabbit 594 (Invitrogen, Cat # A-21207; used 1:1000 in blocking solution). Sections were then washed twice for three minutes then mounted onto slides. All blocking, primary, and secondary antibody steps took place in a humidified chamber. Permeabilization, washes, and antigen retrieval were performed in plastic Coplin jars.

### EdU labeling

EdU labeling was performed either in isolation or following immunofluorescent labeling for other targets. Sections were incubated with the EdU labeling solution for thirty minutes or fifteen minutes at room temperature, washed three times for three minutes, then mounted with ProLong Gold with DAPI. EdU labeling solution: For 1mL, add 40uL of 100mM CuSO_4_ (250mg CuSO_4_ in 10mL 1x TBS), 1uL of 2mM Sulto-Cynine3 azide (Lumiprobe cat# D1330), 100uL of 1M Ascorbic acid sodium salt (20 mg NaAcs (MP Biomedicals cat# 102890) in 100uL 1X TBS) to 860uL 1x TBS.

### TdT-mediated dUTP-X nick end labeling (TUNEL)

The TUNEL assay was performed on paraffin-embedded sections and fifty-micron-thick free-floating frozen sections according to the manufacturer’s instructions but with the time of some steps lengthened to ensure good penetration of the reagents into the tissue (ApopTag® Fluorescein In Situ Apoptosis Detection Kit, Millipore-Sigma, Cat # S7110). Sections were post-fixed for ten minutes at room temperature in 2:1 ethanol:acetic acid, then washed twice for ten minutes with 1x PBS. Equilibration buffer was applied to the sections for three minutes followed by the working strength TdT enzyme for 1.5 hours at 37°C in a humidified chamber. Following the TdT enzyme incubation, stop/wash buffer was applied and incubated at room temperature for fifteen minutes, then the sections were washed three times for three minutes 1x PBS. Working strength anti-digoxigenin conjugate was applied to the sections for one hour at room temperature. Sections were then washed three times for three minutes. Immunofluorescent was then performed as described above.

### Imaging

Ten-micron-thick sections were imaged using a Nikon Eclipse Ti widefield fluorescence microscope using Nikon Plan Fluor 20× (NA: 0.5) or 40× (Oil, NA: 1.3) lenses. An X-Cite 120 LED (Excelitas Technologies) was used. Fifty-micron-thick sections and the in vitro bead engulfment assay were imaged at the Vanderbilt University Cell Imaging Shared Resource using a Zeiss LSM 880 confocal laser scanning microscope. The excitation wavelengths used were 405nm (DAPI), 488 (fluorescein), 561 (Alexa Fluor 594), and 633 (Alexa Fluor 647). The objective lens used was 40x/1.30 C Plan-Apochromat Oil, WD=0.22mm. For the in vivo engulfment analysis (TUNEL and IBA1 co-labeling), the z-step size/voxel depth was 0.427um. For the in vitro bead engulfment, the z-step size/voxel depth was 0.3um. Image analysis and quantification was performed in FIJI (version 1.53r 21 April 2022).

### Quantification

#### In vitro bead engulfment assay

The values collected for this assay allowed the calculation of the percentage of cells performing engulfment and the phagocytic index, defined as (the number of engulfed beads/the total number of cells) x (the number of engulfing/the total number of cells) x 100. The number of engulfed beads was determined using orthogonal views; a bead was considered engulfed if more than ninety percent of the bead surface was found inside the engulfing cell.

#### In vivo analyses

ROIs were traced by hand using the areas of dense DAPI staining as an indicator of the V-SVZ; the lateral ventricles and choroid plexus were excluded from the ROIs, the latter due to high autofluorescence and non-specific staining. The regions outside the ROI were removed using the “Clear outside” command in the Edit dropdown menu to facilitate post-processing and analysis. The sections and thus, the ROIs, fell within the regions described by Merkle et al. as ii/iii (pertaining to their location along the rostro-caudal axis). The dorsal and ventral regions of the V-SVZ on both sides of the brain were analyzed; for simplicity, only dorsal V-SVZ images are shown. Supplementary Figure 6 provides a diagram indicating the location of the images presented in the figures. For EdU and MASH1 labeling, cells were counted as either stained or unstained.

#### TUNEL

Once the V-SVZ ROI was isolated and the regions outside were cleared, several TUNEL^+^ puncta of various sizes were randomly selected and measured using the Line function in the Fiji Toolbar to assess the minimum and maximum punctum size to be included in the analysis. The channels were separated, allowing the TUNEL channel to be thresholded using the Max Entropy algorithm available in the dropdown menu of the Threshold window in Fiji. Once thresholded, the TUNEL^+^ puncta were counted using the Analyze Particles command, using the previously calculated acceptable punctum size. Circularity was left at the default setting. Quantification of puncta in the cortex was performed in the same manner expect the ROI was a large square centered over the S1 region, excluding the corpus callosum, layer I of the cortex, and the meninges. For the fifty-micron-thick sections, the ROIs were determined as described above and the experimenter proceeded through each optical section, identifying and marking ROIs for each TUNEL^+^ cell. A TUNEL^+^ cell was considered engulfed if it was found within in the IBA1-stained area of a microglial cell, either in a process (“ball-and-chain” configuration) or in the soma.

#### IBA1

ROI selection and thresholding for IBA1^+^ area were performed as described above for TUNEL. To measure stained area, the option to limit area measurement to thresholded area was selected in the Set Measurements window found in the Analyze dropdown menu in the Fiji Toolbar. To measure microglia cell number, per cell area, and cell shape using fifty-micron-thick sections, maximum intensity projections (Max IP) were created from the z-stack. For the latter two analyses, the perimeters of fifteen randomly selected microglia were traced and measured, as was the area. The circularity index was defined as 4π[area]/[perimeter]^2^ by Heindl et al.^36^

### Multiplex cytokine analysis

The right and left dorsal V-SVZ were collected from P7 pups according to regional delineations established previously, specifically regions ii/iiiC and ii/iiiD^72^. Each sample contained tissue pooled from 3-6 pups. Tissue was weighed to determine the appropriate volume of lysis buffer (20mM Tris-HCl, 250mM NaCl, 0.05% Tween-20, cOmplete™ Mini Protease Inhibitor Cocktail EASYpacks tablet (1 tablet for 10mL of lysis buffer) (Roche, Cat # 04693124001)) and manually homogenized, then centrifuged at 14K x g for ten minutes at 4°C. The supernatant was removed into a new tube and stored at −80°C until analysis. Analysis of six analytes (interferon γ (IFNγ), interleukin 1β (IL-1β), IL-6, IL-10, IL-17, and tumor necrosis factor α (TNFα)) was performed using the Milliplex Mouse Cytokine Magnetic Kit (EMD Millipore, Cat # MCYTOMAG-70K) and Luminex MAGPIX instrument at the Vanderbilt Hormone Assay and Analytical Service Core. The lysis buffer was used as the matrix solution in the background, standard curve, and control wells.

### Intracerebral injections of interleukin-1 receptor antagonist (IL-1RA)

Intracerebral injections were performed using custom-made clay head molds as described previously^73^. At P4 and weighing 3 grams, pups were euthanized via hypothermia followed by a lethal overdose of 100uL of 2.5% Avertin. Euthanized animals were placed in 3% agarose such that only the head and shoulders were submerged, and the agarose was left to solidify. The pups were carefully removed from the agarose, and the cavity left behind was filled with dental repair resin and allowed to cure for 24-72 hours at room temperature in a well-ventilated area. The resin casts were used as models around which to mold Crayola Air-Dry modeling clay; the cast was placed such that the dorsal aspect of the head was oriented up and was visible. The clay was allowed to harden, and the mold was affixed to the top of an inverted Petri dish for stability once under the custom-built injecting rig (kindly provided by Dr. Ihrie). Prior to the injection, pups were placed in a protective nitrile sleeve, anesthetized on ice for 5-10 minutes, and then immobilized in the clay head molds using tape; the skin covering the cranium was held taut by transparent tape. A beveled glass needle was pre-loaded with mineral oil and the recombinant human IL-1 receptor inhibitor (IL-1RA, R&D Systems, Cat # 280-RA, resuspended in 0.1% BSA), then positioned in the injecting rig at a 90° angle relative to the horizontal plane of the Petri dish and head mold. The needle was centered along the midline between the eyes and the position was calibrated as (x = 0mm, y = 0mm, z = 0mm); the injection site was 3mm caudal and 0.5mm lateral to that starting site, and 3mm down into the brain parenchyma. Pups received 1uL of 1ug/uL IL-1RA or 0.1% BSA at these coordinates in the right hemisphere only. To differentiate between the pups that had already received an injection, prior to arousal from anesthesia, the paw was tattooed; each pup had a different combination of tattoos on their fore and rear paws and tail. The pups were placed on a Deltaphase® isothermal pad and observed until the suckling and righting reflexes returned, then they were placed in the home cage with the dam and littermates. Pups were observed daily for the next three days for abnormalities. At P7, EdU was administered one hour prior to euthanasia and processing as described above.

### Tamoxifen induction of microglia-specific Jedi knockout

100mg of Tamoxifen (Sigma-Aldrich, Cat # T5648-1G) was resuspended in 1mL pure ethanol and 9mL peanut oil (Sigma-Aldrich, Cat # P2144-250ML) and administered to dams, twice daily, when the pups were P1 to P3. Intraperitoneal injections (100ul, containing 1mg Tamoxifen) were performed immediately rostral to the inguinal white adipose tissue, alternating sides between injections.

### Magnetic-activated cell sorting (MACS)

Cortical hemispheres (containing the V-SVZ) were collected as described for above (see Multiplex cytokine analysis) from control and microglia-specific Jedi KO mice. The overlaying cortex was also collected and placed in a separate tube. The samples were dissociated using the Papain Dissociation System (see ‘Primary microglia cultures’), and microglia were then isolated from the cell suspension by Magnetic Activated Cell Sorting using Cd11b microbeads per manufacturer instructions with MS columns and MiniMACs separator (Miltenyi Biotec, Cat # 130-093-634).

### Assessment of Tamoxifen-induced removal of Jedi in microglia by qPCR

RNA was extracted from MACS-isolated whole-cortex microglia using the RNeasy Micro Kit per manufacturer instructions (Qiagen, Cat # 74004) and quantified on a NanoDrop 2000 (ThermoFisher Scientific, Cat # ND-2000). cDNA was prepared from 50ng RNA using the SuperScript III First-Strand synthesis system (Invitrogen, Cat # 18080051). Quantitative PCR for Jedi and Gapdh was performed using Sso Universal SYBR Green Supermix (BioRad, Cat # 1725271) on the CFX Connect Real-Time System (BioRad, Cat t# 1855201). The primers are listed in Table 2.

**Table 2.**
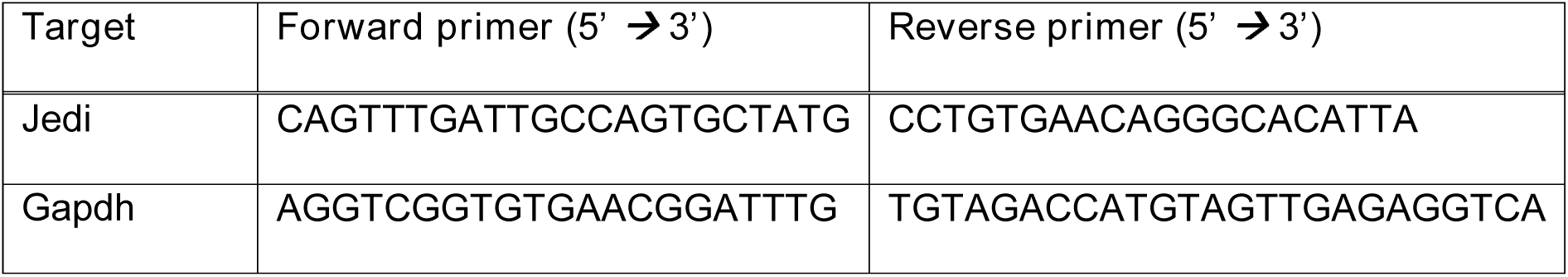
Quantitative PCR primer sequences

### Bulk RNA sequencing

#### Library preparation

RNA was isolated as described above. All RNA-seq experiments were performed in biological replicates. RNA was submitted to Vanderbilt University Medical Center VANTAGE Core for poly-A enrichment-based library preparation and sequencing on the Illumina NovaSeq (PE-100).

#### Data analysis

Pre-processed reads were aligned to the mouse genome (mm10, downloaded from UCSC) using TopHat (v2.0.11)^74^ and differential gene expression was determined using DESeq2 (v. 1.34.0) as previously described^75^. Sequencing run was included in DESeq assay design to control for batch effect. Genes were considered differentially expressed if they had an adjusted p-value < 0.05 and a fold change ≥ |2|. Differentially expressed genes were used for gene ontology analysis using clusterProfiler (v. 4.2.2)^76^.

### Liquid chromatograph-mass spectrometry

Samples were collected from the dorsal V-SVZ of P7 WT and JKO pups, snap frozen in liquid nitrogen, and stored at −80°C until LC-MS analysis.

#### Materials

HPLC-grade methanol, acetonitrile, isopropanol and formic acid used for sample purification and LC-MS/MS analysis were JT Baker-brand (ThermoFisher Scientific, Waltham, MA). Authentic lipid standards of 6-keto-PGF_1α_, PGE_2_, PGD_2_, PGF_2α_, PGJ_2_, 15-hydroxyeicosanoic acid (15-HETE), 2-arachidonoylglycerol (2-AG), anandamide (AEA), arachidonic acid (AA), oloeylglycerol (OG), and oleoylethanolamide (OEA) were purchased from Cayman Chemicals (Ann Arbor, MI). 16:0 lysophosphatidylcholine (LPC) and 17:0 LPC were purchased from Avanti Polar Lipids (Alabster, AL). The following deuterated internal standards were purchased from Cayman: 6-keto-PGF_1α_-d4, PGF_2α_-d4, PGE_2_-d4, PGD_2_-d4, PGJ_2_-d4, 15-HETE-d8, 2-AG-d5, AEA-d4, AA-d8, and OEA-d4. OG-d5 was purchased from US Biological (Salem, MA).

All LC-MS/MS analysis was performed on a Shimadzu Nexera system in-line with a SCIEX 6500 QTrap; except the LPC analysis, which was performed on a Shimadzu LC system in-line with a SCIEX 3200 QTrap mass spectrometer. The 6500 QTrap was equipped with a TurboV Ionspray source and operated in positive and negative ion modes. The 3200 QTrap was equipped with an ionspray source and operated in positive ion mode. For both LC-MS systems, SCIEX Analyst software (ver 1.6.2) was used to control the instruments and acquire and process the data.

#### Sample preparation

Vials containing collected brain samples were removed from −80°C storage and placed on dry ice. 0.5 mL of methanolic homogenization solution containing the deuterated internal standards were added to each vial and the vial was sonicated in a bath sonicator filled with ice water. After about 1 minute of sonication, the samples were pulverized completely. The samples were then transferred to glass test tubes and centrifuged at ∼3,000 rcf for ten minutes at 4°C. Roughly 0.4 mL of the supernatant was removed to clean test tubes and evaporated to dryness under nitrogen and either reconstituted for LC-MS analysis or capped and stored at −20°C for later analysis.

#### Lipid analysis

Immediately prior to LC-MS analysis, the samples were reconstituted in 70µL MeOH and 40µL H_2_O, vortexed and transferred to autosampler vials. The samples were analyzed on the above-referenced LC/MS systems. For LPC analysis, the analytes were chromatographed on an Phenomenex C18 column (5.0 x 0.21 cm; 3.0 μm) which was held at 43°C. A gradient elution profile was applied to each sample, specifically; an initial hold for 0.30 min followed by an increase in %B from 15% (initial conditions) to 90% over 5.0 min, and a hold at 90% for 0.5 min. The column was subsequently returned to initial conditions for 1.5 min prior to the next injection. The flow rate was 300 μL/min and component A was 1:1 methanol:water with 5 mM ammonium acetate and 0.1% formic acid while component B was 2:1:1 isopropanol:acetonitrile:methanol with 5 mM ammonium acetate and 0.1% formic acid.

For all other lipids, the analytes were chromatographed on an Acquity UPLC BEH C18 reversed-phase column (5.0 x 0.21 cm; 1.7 μm) which was held at 43°C. A gradient elution profile was applied to each sample, specifically; an initial hold for 0.25 min followed by an increase in %B from 35% (initial conditions) to 99% over 4.00 min, and a hold at 99% for 0.8 min. Then the column was returned to initial conditions for 1.2 min prior to the next injection. The flow rate was 330 μL/min and component A was water with 0.05% formic acid while component B was 3:1 acetonitrile:methanol with 0.1% formic acid.

All analytes were detected via selected reaction monitoring (SRM) in both negative and positive ion mode. The SRM transition (m/z), collision energy (C.E. - volts) and polarity are given for each analyte in Table 4. Italicized values in parentheses indicate that value for the deuterated internal standard. Analytes were quantitated by stable isotope dilution against their deuterated internal standard, except 16:0 LPC, which was quantitated against 17:0 LPC. Data were normalized to picomoles of 16:0 LPC per gram of tissue.

**Table 4.** SRM Reactions of Lipid Analytes

| **Analyte** | **Q1 (m/z)** | **Q3 (m/z)** | **C. E.** | **Polarity** |
| --- | --- | --- | --- | --- |
| **6-keto-PGF<sub>2α</sub>** | 369.2 (373.2) | 163.1 (167.1) | -32 | - |
| **15-HETE** | 319.2 (327.2) | 169.1 (173.1) | -20 | - |
| **PGF<sub>2α</sub>** | 353.2 (357.2) | 193.1 (197.1) | -34 | - |
| **PGE<sub>2</sub> & D<sub>2</sub>** | 351.2 (355.2) | 271.1 (275.1) | -23 | - |
| **PGJ<sub>2</sub> & 15-d-PGD<sub>2</sub>** | 333.2 (337.2) | 271.1 (275.1) | -22 | - |
| **15-d-PGJ<sub>2</sub>** | 315.2 (319.2) | 271.1 (275.1) | -25 | - |
| **2-AG** | 379.2 (384.2) | 287.2 (287.2) | 29 | + |
| **AEA** | 348.2 (352.2) | 62.0 (66.0) | 24 | + |
| **AA** | 303.1 (311.1) | 259.0 (267.0) | -15 | - |
| **OG** | 357.2 (362.2) | 265.2 (265.2) | 23 | + |
| **OEA** | 326.2 (330.2) | 62.0 (66.0) | 36 | + |
| **16:0 LPC** | 496.4 | 184.2 | 35 | + |
| **17:0 LPC** | 510.4 | 184.2 | 35 | + |

## Acknowledgements

We thank M. Tugend and F. E. Hickman for their work at the early stage of the project. We thank A. Sierra for her thoughtful discussions and guidance.

## Supplementary Figures

**Supplementary Figure 1.**
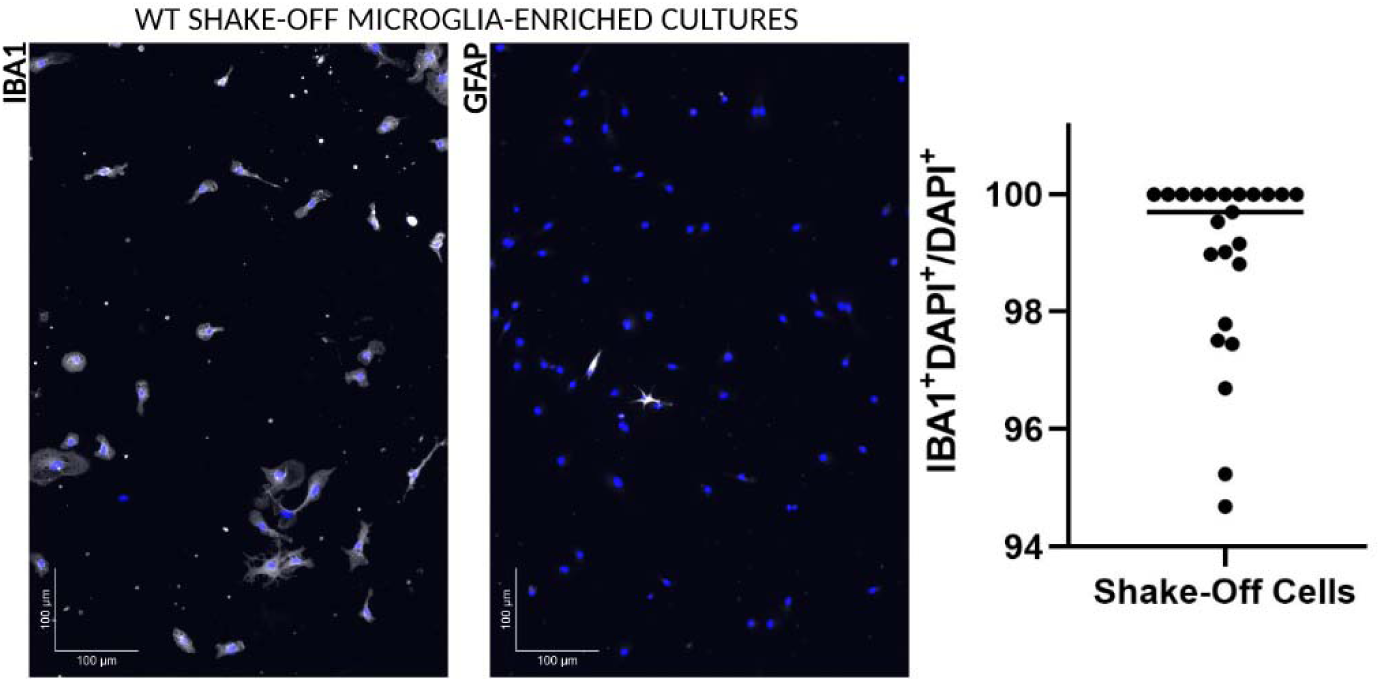
Shake-off microglia culture is highly pure. Left: DAPI (blue) and IBA1 (white) staining in wild-type (WT) microglia cultures (days in vitro 18 (DIV18)). Middle: DAPI (blue) and GFAP (white, astrocyte marker) staining in WT microglia cultures. (Due to antibody incompatibilities, these stains could not be performed together and thus, were done in neighboring wells of a chamber-slide, but all cells were derived from the same mixed glia culture.) Right: Quantification of IBA1+ cells in WT microglia cultures shows that 98.89% of cultured cells in the chamber stained for IBA1 are IBA1+. In the chamber stained for GFAP, GFAP+ astrocytes made up 1.14% of cultured cells.

**Supplementary Figure 2.**
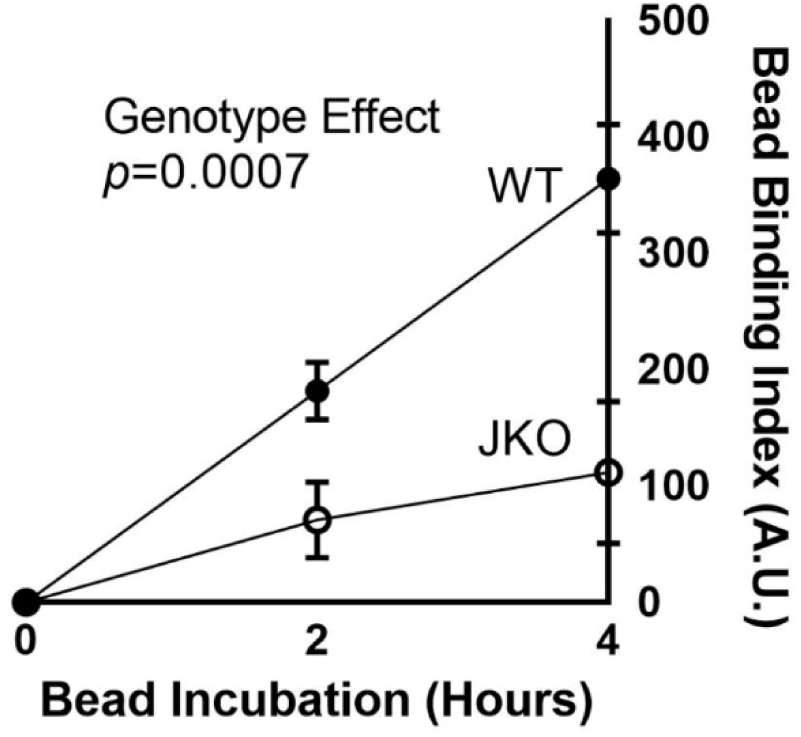
Bead binding index is reduced in JKO microglia in vitro. Quantification of the ‘Bead Binding Index’ ((the number of attached beads/the total number of cells) x (the number of engulfing/the total number of cells) x 100) in wild-type (WT) and Jedi knockout (JKO) microglia cultures. Data points are the average of three independent cultures and engulfment assays. Error bars represent the standard error of the mean. Arbitrary units, A.U.

**Supplementary Figure 3.**
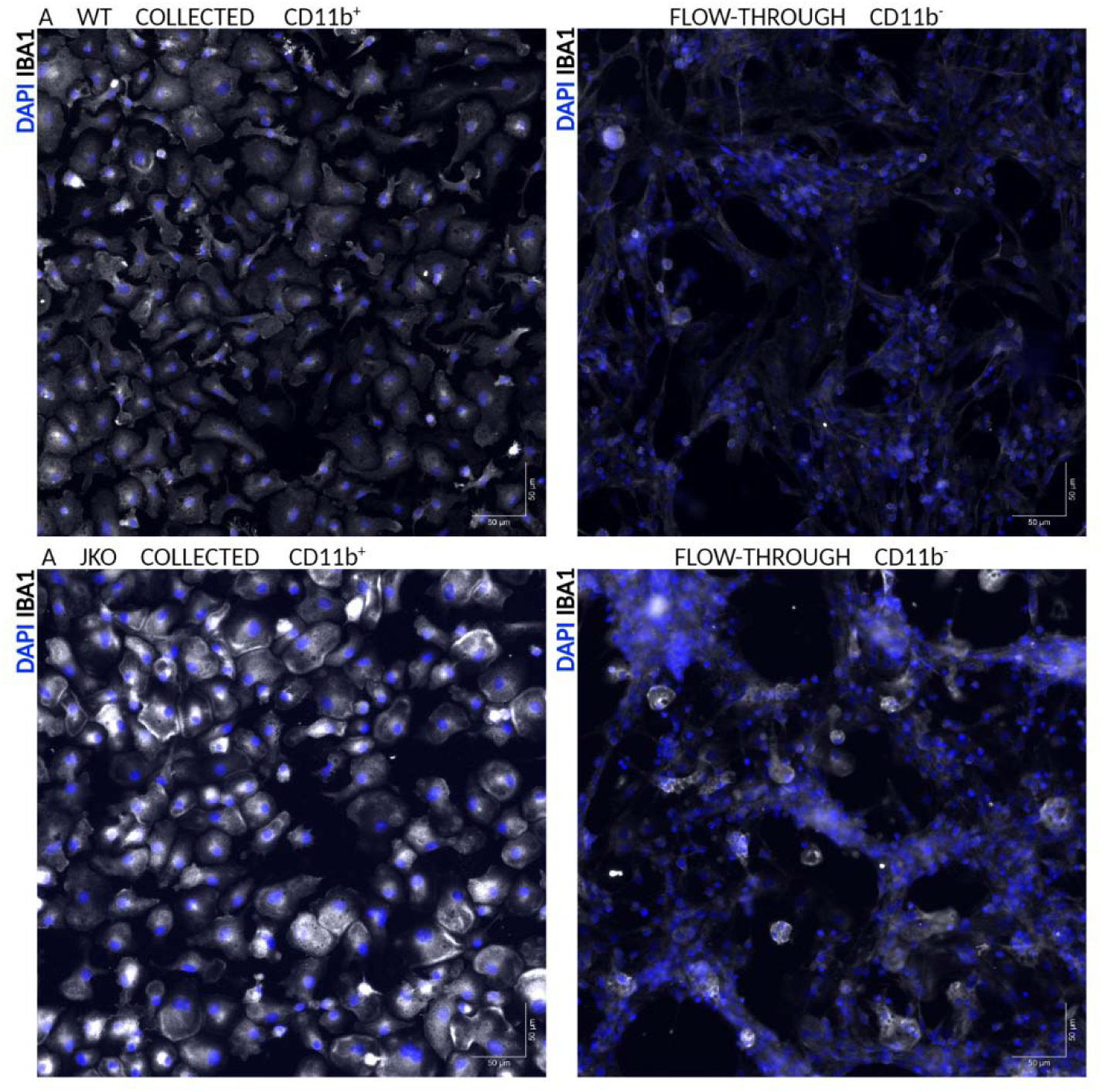
Immunomagnetically sorted microglia are highly pure. S3A. Left: Wild-type (WT) microglia (IBA1, white) following magnetic cell sorting based on CD11b expression. Right: Flow-through fraction that did not express CD11b. Cell nuclei are labeled with DAPI (blue). S3B. Left: Jedi knockout (JKO) microglia (IBA1, white) following magnetic cell sorting based on CD11b expression. Right: Flow-through fraction that did not express CD11b. Cell nuclei are labeled with DAPI (blue).

**Supplementary Figure 4.**
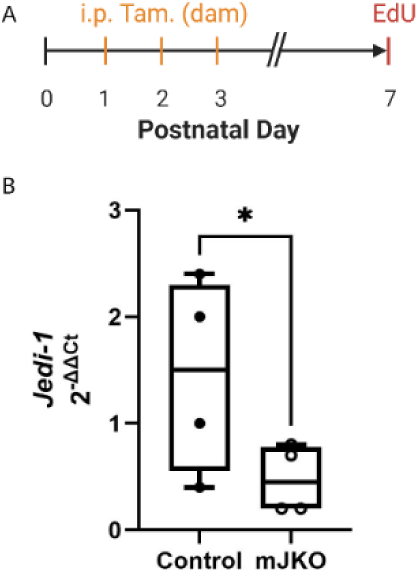
Tamoxifen-induced removal of Jedi in microglia-specific knockout line. S4A. Timeline of Tamoxifen (Tam)-induced deletion of Jedi in Cx3cr1CreERT2/+;Jedifl/fl microglia-specific JKO (mJKO). Tam was administered intraperitoneally (i.p.) to the post-partum dam twice daily from postnatal day 1 (P1) to P3. Then at P7, EdU (5-ethynyl-2’-deoxyuridine) was delivered directly to the pups to mark proliferating cells one hour before euthanasia. Littermates (also received Tam) with the following genotypes were used as controls (Cx3cr1+/+, Cx3cr1CreERT2/+;Jedifl/+, or Cx3cr1CreERT2/+;Jedi+/+). S4B. Quantitative polymerase chain reaction in immunomagnetically sorted whole-cortex microglia shows successful reduction of Jedi-1 (‘Jedi’) messenger RNA in mJKO pups compared to control littermates.

**Supplementary Figure 5.**
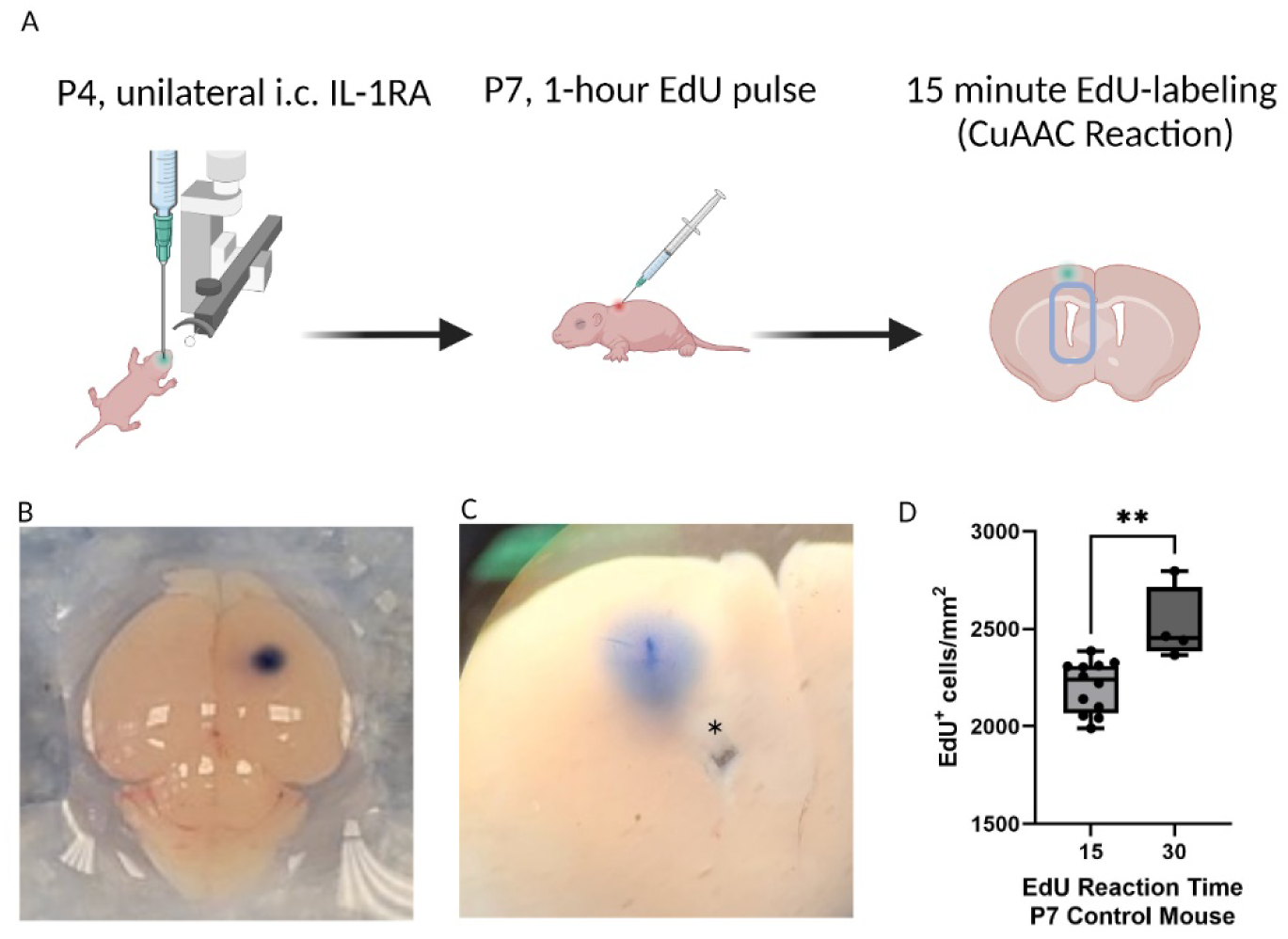
IL-1R antagonism rescues neurogenesis in the JKO V-SVZ. S5A. Workflow schematic of interleukin-1 receptor antagonist (IL-1RA) administration: Left: Postnatal day 4 (P4) intracerebral (i.c.) injection of IL-1RA in the right hemisphere. Middle: Subcutaneous injection of EdU (5-ethynyl-2’-deoxyuridine) at P7 to mark proliferating cells one hour before euthanasia. Right: The EdU labeling reaction (CuAAC, Cu (I)-catalyzed azide-alkyne cycloaddition) was performed for fifteen minutes rather than thirty minutes, the latter of which was used in preceding experiments. S5B. Dorsal view of injection site on P4 mouse brain. Trypan blue was injected to visualize the injection site. S5C. Trypan blue indicates the injection site in this anterior view of a coronal slice of the P4 brain (top of the image is dorsal, right side of the image is medial). Asterisk indicates the lateral ventricle. S5D. Quantification of EdU+ cells in the ventricular-subventricular zone (V-SVZ) of a P7 wild-type pup following a 15-minute or 30-minute CuAAC reaction. Unpaired one-tailed t test, ** p=0.0012.

**Supplementary Figure 6.**
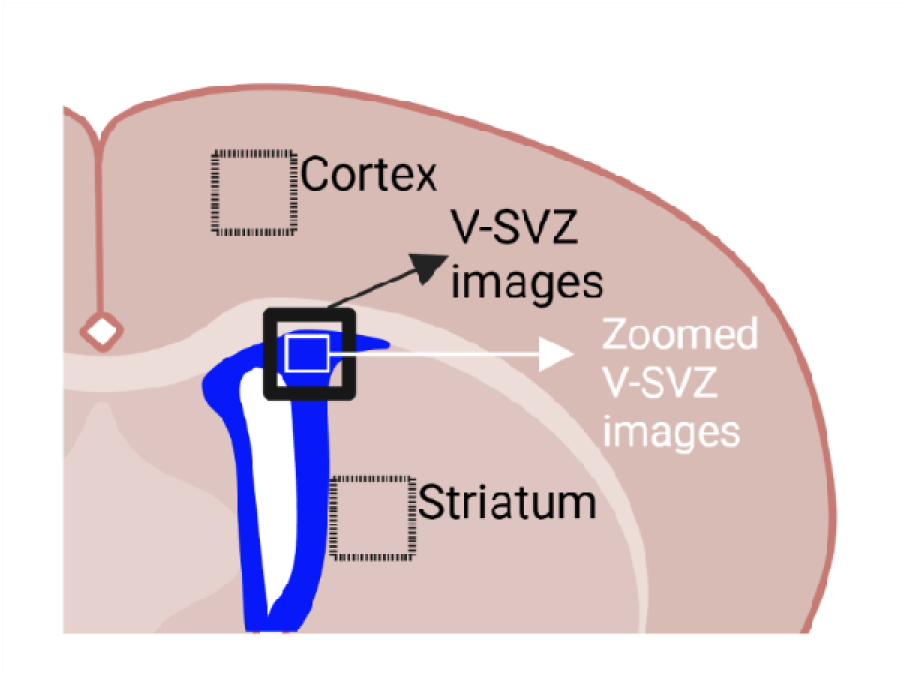
Diagram indicating brain regions presented in the figures. A coronal view of the mouse brain indicates the regions presented in the figures. Data were collected from both the dorsal and ventral V-SVZ on both sides of the brain. For simplicity, only images of the dorsal V-SVZ are presented in the figures. The boxes with dotted lines correspond to the cortex and striatum (in Figure 1B and 1C, and Figure 3E and 3F). The box with a solid black line corresponds to where all other images for the figures were taken, with the following exceptions: (1) Figure 10F, 10G, 10I, 10H, 11A, and 11B are in the opposite orientation relative to most other images because they happened to have been acquired on the other side of the brain slices being imaged. (2) Images in the following panels correspond to the small white box within the V-SVZ: Figure 5A, 5B, 5D, and 5E; Figure 10a and 10b (for both MASH1 and EdU), and 10F, 10G, 10I, and 10J.

